# Enteroendocrine cell lineages that differentially control feeding and gut motility

**DOI:** 10.1101/2022.03.18.484842

**Authors:** Marito Hayashi, Judith A. Kaye, Ella R. Douglas, Narendra R. Joshi, Fiona Gribble, Frank Reimann, Stephen D. Liberles

## Abstract

Enteroendocrine cells are specialized sensory cells of the gut-brain axis that are sparsely distributed along the intestinal epithelium. The functions of enteroendocrine cells have classically been inferred by the gut hormones they release. However, individual enteroendocrine cells typically produce multiple, sometimes apparently opposing, gut hormones in combination, and some gut hormones are also produced elsewhere in the body. Here, we developed approaches involving intersectional genetics to enable selective access to enteroendocrine cells *in vivo*. We constructed *Villin1-p2a-FlpO* knock-in mice to restrict reporter expression to intestinal epithelium and through combined use of Cre and Flp alleles, effectively targeted major transcriptome-defined enteroendocrine cell lineages that produce serotonin, glucagon-like peptide 1, cholecystokinin, somatostatin, or glucose insulinotropic peptide. Chemogenetic activation of different enteroendocrine cell types variably impacted feeding behavior, nausea-associated behaviors (conditioned flavor avoidance), and gut motility. Defining the physiological roles of different enteroendocrine cell types provides an essential framework for understanding sensory biology of the intestine.

## INTRODUCTION

The gut-brain axis plays a critical role in animal physiology and behavior. Sensory pathways from the gut relay information about ingested nutrients, meal-induced tissue distension, osmolarity changes in the intestinal lumen, and cellular damage from toxins (Bai et al., 2019; Brookes et al., 2013; Prescott and Liberles, 2022; Richards et al., 2021; Williams et al., 2016). Responding neural circuits evoke sensations like satiety and nausea, coordinate digestion across organs, shift systemic metabolism and energy utilization, and provide positive and negative reinforcement signals that guide future consumption of safe, energy-rich foods (Andermann and Lowell, 2017; Sternson and Eiselt, 2017; Zimmerman and Knight, 2020). Moreover, manipulations of the gut-brain axis have been harnessed clinically through gut hormone receptor agonism or bariatric surgery to provide powerful therapeutic approaches for obesity and diabetes intervention (Richards *et al.*, 2021; Seeley et al., 2015).

Enteroendocrine cells are first-order chemosensory cells of the gut-brain axis, and are sparsely distributed along the gastrointestinal tract (Gribble and Reimann, 2019). Like taste cells, enteroendocrine cells are epithelial cells with neuron-like features, as they are electrically excitable, release vesicles upon elevation of intracellular calcium, and form synaptic connections with second-order neurons through specialized extrusions called neuropods (Bohorquez et al., 2015; Reimann et al., 2012). Single-cell RNA sequencing approaches revealed a diversity of enteroendocrine cell types that produce different gut hormones (Beumer et al., 2018; Gehart et al., 2019; Haber et al., 2017). Superimposing cell birthdate on the enteroendocrine cell atlas through an elegant genetically encoded fluorescent clock revealed five major enteroendocrine cell lineages defined by expression of either glucose insulinotropic peptide (GIP), ghrelin, serotonin (called enterochromaffin cells), somatostatin, or a combination of glucagon-like peptide 1 (GLP1), cholecystokinin (CCK), and/or neurotensin (Gehart *et al.*, 2019).

Enteroendocrine cell-derived gut hormones evoke a variety of physiological effects (Drucker, 2016). GLP1 and CCK are satiety hormones released following nutrient intake, ghrelin is an appetite-promoting hormone whose release is suppressed by nutrients, and serotonin is released by non-nutritive signals like irritants, force, catecholamines, and small chain fatty acids. Sugar-induced release of GIP and GLP1 causes the incretin effect which rapidly promotes insulin release and lowers blood glucose (Holst et al., 2009). CCK, serotonin, and other gut hormones additionally regulate a variety of digestive functions including gut motility, gastric emptying, gastric acidification, absorption, gallbladder contraction, and exocrine pancreas secretion.

The functions of individual enteroendocrine cell types could in some cases be inferred by summing the actions of their expressed hormones. For example, chemogenetic activation of enteroendocrine cells in the distal colon which express insulin-like peptide 5 triggers a multi-pronged physiological response that includes appetite suppression through a peptide YY (PYY) receptor, improved glucose tolerance through GLP1, and defecation indirectly through the serotonin receptor HTR3A (Lewis et al., 2020). However, a challenge in generalizing this approach is that some enteroendocrine cells release hormones with apparently opposing functions (Gehart *et al.*, 2019; Haber *et al.*, 2017), and moreover, many gut hormones are also produced by other cell types in the body (Lee and Soltesz, 2011; Okaty et al., 2019). To overcome these challenges, we developed approaches involving intersectional genetics to obtain highly selective access to major transcriptome-defined enteroendocrine cell lineages. Chemogenetic activation of each of these enteroendocrine cell types produced variable effects on gut physiology and behavior. Obtaining a holistic model for enteroendocrine cell function provides a critical framework for understanding the neuronal and cellular logic underlying gut-brain communication.

## RESULTS AND DISCUSSION

### Selective access to enteroendocrine cells *in vivo* through intersectional genetics

We first sought to identify genetic tools which broadly and selectively mark enteroendocrine cells. Transcription factors such as ATOH1, Neurogenin3, and NeuroD1 are expressed in enteroendocrine cell progenitors and/or precursors and act in early stages of enteroendocrine cell development (Li et al., 2011). We obtained *Atoh1-Cre* (both knock-in and transgenic lines), *Neurog3-Cre*, and *Neurod1-Cre* mice and crossed them to mice containing a Cre-dependent tdTomato reporter (*lsl-tdTomato*). *Neurog3-Cre* and *Neurod1-Cre* lines labeled a sparse population of intestinal epithelial cells characteristic of enteroendocrine cells, although the *Neurog3-Cre* line additionally labeled other cells in intestinal crypts and in occasional mice produced broad labeling of intestinal epithelium; neither *Atoh1-Cre* line tested displayed selective labeling of enteroendocrine cells (Figure 1 – figure supplement 1A) (Schonhoff et al., 2004). Two-color analysis of tdTomato and gut hormone expression verified tdTomato localization in enteroendocrine cells of *Neurod1-Cre*; *lsl-tdTomato* mice, consistent with prior findings (Figure 1 – figure supplement 1B) (Li et al., 2012). Single cell RNA sequencing of tdTomato-positive cells obtained from these mice (see below) also verified selective enteroendocrine cell labeling.

*Neurod1-Cre* mice provide broad, indelible, and selective marking of enteroendocrine cells within the intestine, but NeuroD1 is also expressed in a variety of other tissues, including the brain, retina, pancreas, peripheral neurons, and enteric neurons (Figure 1B, C) (Cho and Tsai, 2004; Li *et al.*, 2011). Knockout of NeuroD1 is lethal, causing severe deficits in neuron birth and survival, as well as in development of pancreatic islets and enteroendocrine cells (Gao et al., 2009; Naya et al., 1997). We employed an intersectional genetic strategy of combining Cre and Flp recombinases to limit effector gene expression to enteroendocrine cells. *Villin1* (*Vil1*) is expressed with high selectivity in the lower gastrointestinal tract (el Marjou et al., 2004; Maunoury et al., 1992), so we generated a knock-in mouse allele (*Vil1-p2a-FlpO*) that drives FlpO recombinase expression from the endogenous *Vil1* locus. *Vil1-p2a-FlpO* mice displayed expression of a Flp-dependent *Gfp* allele in epithelial cells throughout the entire length of the intestine with striking specificity (Figure 1A, Figure 1 – figure supplement 1C). Reporter expression was not observed in most other tissues examined, including most brain regions, spinal cord, peripheral ganglia, and enteric neurons; rare GFP-expressing cells were noted in taste papillae, epiglottis, pancreas, liver, and thalamus (Figure 1C, Figure 1 – figure supplement 1C, E) (Hofer and Drenckhahn, 1999; Madison et al., 2002; Rutlin et al., 2020). Combining *Neurod1-Cre* and *Vil1-p2a-FlpO* alleles yielded highly selective expression of an intersectional reporter gene encoding tdTomato (*inter-tdTomato*) in enteroendocrine cells, with only occasional cells observed in pancreas, and no detectable expression in other cell types labeled by either allele alone (Figure 1C, Figure 1 – figure supplement 1D, E).

**Figure 1.**
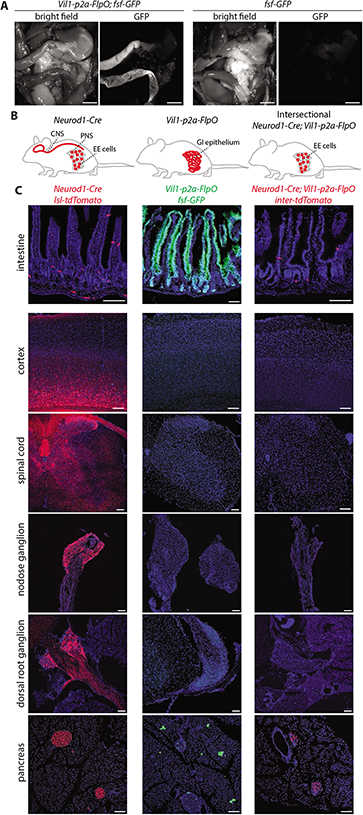
Establishing intersectional tools for genetic access to enteroendocrine cells *in vivo*. (A) Bright field microscopy and native GFP fluorescence microscopy of intestinal tissue from *Vil1-p2a-FlpO; fsf-Gfp* mice (left) and *fsf-Gfp* mice (right), scale bar: 5 mm. (B) Cartoon depicting intersectional genetic strategy to access enteroendocrine cells. (C) Native reporter fluorescence in cryosections (20 μm, except 50 μm for cortex, spinal cord) of fixed tissues indicated from *Neurod1-Cre; lsl-tdTomato* mice (left), *Vil1-p2a-FlpO; fsf-Gfp* mice (middle), and *Neurod1-Cre; Vil1-p2a-FlpO; inter-tdTomato* mice (right), scale bar: 100 μm, intestine sections from duodenum (middle) or jejunum (left, right). See Figure 1 - figure supplement 1.

### Charting enteroendocrine cell diversity and gene expression

Our general goal was to use intersectional genetics to access subtypes of enteroendocrine cells that express different gut hormones. We first used single cell RNA sequencing approaches to measure the extent of enteroendocrine cell diversity, to compare findings with existing enteroendocrine cell atlases, and to establish a foundation for genetic experiments. Enteroendocrine cells represent <1% of gut epithelial cells, so we used genetic markers for enrichment. NeuroD1 is expressed early in the enteroendocrine cell lineage, and we observed by two-color expression analysis that *Neurod1-Cre* mice target at least several enteroendocrine cell types (Figure 1 – figure supplement 1B). Since prior enteroendocrine cell atlases were derived from cells expressing an earlier developmental marker, *Neurog3* (Gehart *et al.*, 2019), we sought to compare the repertoire of enteroendocrine cells captured by *Neurod1-Cre* and *Neurog3-Cre* mice.

tdTomato-positive cells were separately obtained from the intestines (duodenum to ileum) of *Neurod1-Cre; lsl-tdTomato* mice and *Neurog3-Cre; lsl-tdTomato* mice by fluorescence-activated cell sorting (Figure 2 – figure supplement 1A). Using the 10x Genomics platform, mRNA was captured from individual cells, and barcoded single cell cDNA was generated. Single cell cDNA was then sequenced and unsupervised clustering analysis was performed using the Seurat pipeline (Hafemeister and Satija, 2019; Stuart et al., 2019). Transcriptome data was obtained for 5,856 tdTomato-positive cells from *Neurog3-Cre; lsl-tdTomato* mice and 1,841 tdTomato-positive cells from *Neurod1-Cre; lsl-tdTomato* mice. 25% of *Neurog*3-lineage cells (1454/5856) and 87% of NeuroD1-lineage cells (1595/1841) expressed classical markers for enteroendocrine cells (Figure 2 – figure supplement 1B, C). Moreover, the full diversity of known enteroendocrine cell types was similarly captured by both Cre lines, with *Neurog3-Cre* mice additionally labeling many other cells, including paneth cells, goblet cells, enterocytes, and progenitors (Figure 2 – figure supplement 1C). These findings are consistent with NeuroD1 acting later than Neurogenin3 in the enteroendocrine cell lineage, but prior to cell fate decisions leading to enteroendocrine cell specialization (Jenny et al., 2002).

Since *Neurog3-Cre* and *Neurod1-Cre* mice similarly labeled all known enteroendocrine cell lineages, transcriptome data was computationally integrated for analysis of enteroendocrine cell subtypes. Selective clustering analysis of 3,049 enteroendocrine cells from both mouse lines revealed 10 distinct cell clusters, with one cluster representing putative progenitors (Figure 2A, Table 1). Cell clusters were compared with previously described enteroendocrine cell types based on expression of signature genes encoding hormones and transcriptional regulators (Figure 2A-C) (Gehart *et al.*, 2019). We observed three classes of enterochromaffin cells that similarly express serotonin biosynthesis enzymes (*Tph1*) and associated transcription factors (*Lmx1a*), but differentially produce *Tac1*, *Cart*, *Pyy*, *Ucn3*, and *Gad2* (Figure 2B). Six other cell types preferentially express either *Gip* (K cells), *Cck* (I cells), *Gcg* (GLP1 precursor, L cells), *Nts* (N cells), *Sst* (D cells), and *Ghrl* (X cells), with L, I, and N cells thought to be derived from a common cell lineage (Beumer et al., 2020; Gehart *et al.*, 2019). Strong segregation was observed for some signature genes, such as *Tph1* in enterochromaffin cells and *Sst* in D cells. In other cases, signature hormone genes like *Cck* and *Ghrl* were enriched in particular cell clusters but expression was not absolutely restricted and also observed at lower levels in other cell clusters (Figure 2B). We note that glutamate transporters were not readily detected in our transcriptomic data. Thus, each enteroendocrine cell subtype expresses a hormone repertoire with distinct patterns of enrichment but also sometimes partial overlap.

**Figure 2.**
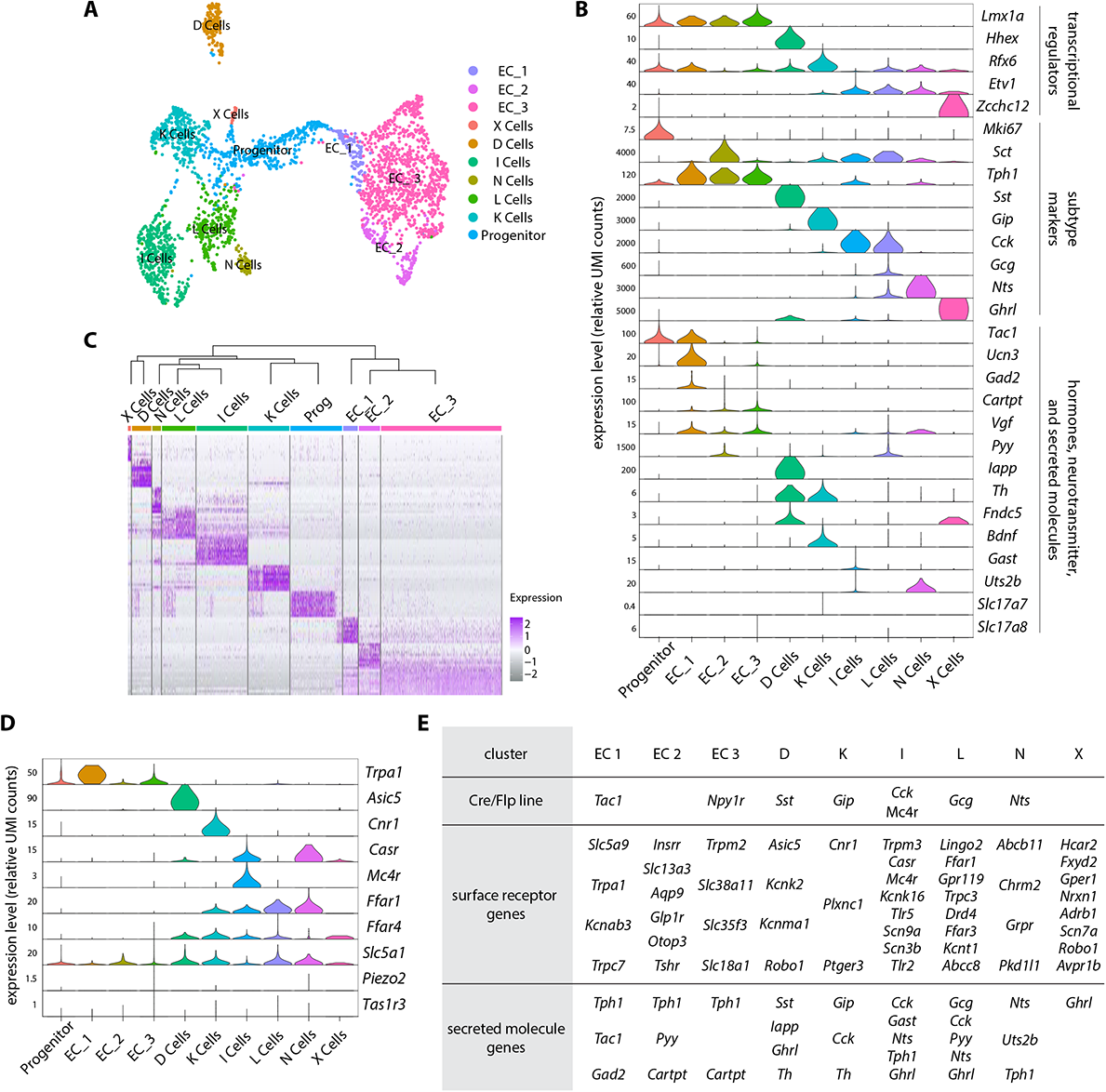
An enteroendocrine cell atlas reveals differential hormone and receptor expression. (A) A uniform manifold approximation and projection (UMAP) plot of enteroendocrine cell transcriptomic data reveals 10 cell clusters. (B) Violin plots showing gene expression across enteroendocrine cell subtypes. (C) Normalized expression of enriched signature genes (see Table 1 for a gene list) in single enteroendocrine cells. The dendrogram (top) depicts the relatedness (quantified by position along the Y-axis) between cell clusters based on gene expression. (D) Violin plots showing expression of cell surface receptor genes across enteroendocrine cell subtypes. (E) For each enteroendocrine cell type, examples of gene loci used for genetic targeting (top), expressed cell surface receptor genes (middle) and expressed hormone and neurotransmitter-related genes (bottom). Genes were selected among the top 30 differentially expressed genes. See Figure 2 - figure supplement 1.

**Table 1.** Signature genes with differential expression across enteroendocrine cell types.

| progenitor | EC 1 | EC 2 | EC 3 | D cells | K cells | I cells | L cells | N cells | X cells |
| --- | --- | --- | --- | --- | --- | --- | --- | --- | --- |
| Gvin-ps1.1 | Atp6v1c2 | Lmo3 | Iqsec1 | Gm30613 | Cnr1 | Pzp | Lingo2 | Rassf9 | Irs4 |
| Myb | Hip1 | Foxq1 | Trpm2 | Hhex | Dlgap2 | Habp2 | Col8a1 | Hs3st2 | Tex36 |
| Kcnn4 | Cidea | Ctse | Dlg2 | Kcnk2 | Sphkap | Crp | Ffar1 | Adgrd1 | Adcy5 |
| Gvin-ps1 | Csgalnact1 | Fbxl7 | Lmx1a | Gm21149 | Slfn10-ps | Cyp2j5 | Naaladl1 | Gm34237 | Olfml2b |
| Kcne3 | Alb | Insrr | Slc38a11 | Sez6l | Pkib | Adamts5 | Gcg | Abcb11 | Olfml3 |
| Cdca7 | Ankrd55 | Hs3st5 | Cdk14 | Iapp | Plcxd3 | Basp1 | Tmem108 | Sncg | Gm11837 |
| Smoc2 | Slc5a9 | Gm15218 | Ccdc3 | Fam151a | Slfn8 | Trpm3 | Tnr | Rgs6 | Tdh |
| Nhs | Fabp3 | Cwh43 | Snap25 | Asic5 | Plcb1 | Galnt13 | Etv1 | Rims3 | Gm26862 |
| BC064078 | Gm45740 | Meis2 | Pde4d | Sptbn2 | Pdzrn3 | Casr | Map2 | Gm47833 | Hcar2 |
| Cftr | Gstk1 | Slc13a3 | Tph1 | Gm7205 | Plxnc1 | Ugt8a | Gm34701 | Samd11 | Plb1 |

Enteroendocrine cells also express various cell surface receptors to detect nutrients, toxins, and other stimuli. For example, enteroendocrine cells detect sugars through the sodium-glucose cotransporter SGLT1 (encoded by the gene *Slc5a1*), with sodium co-transport thought to lead directly to cell depolarization (Gorboulev et al., 2012; Reimann et al., 2008). This mechanism is distinct from sugar detection by taste cells or pancreatic beta cells. Gustatory sensations of sweet (and savory/umami) involve taste cell-mediated detection of sugars (and amino acids) through heterodimeric G Protein-Coupled Receptors termed T1Rs (Yarmolinsky et al., 2009), while pancreatic beta cells respond to sugar through increased metabolic flux, ATP-gated potassium channel closure, and depolarization. Expression of *Slc5a1* was observed in multiple enteroendocrine cell subtypes, and highest in K, L, D, and N cells, while abundant expression of T1Rs was not detected in any enteroendocrine cell type (Figure 2D). These findings are consistent with the ability of taste blind mice lacking T1Rs to develop a preference for sugar-rich foods through SGLT1-mediated post-ingestive signals of the gut-brain axis (Sclafani et al., 2016; Tan et al., 2020). In addition, free fatty acid receptor genes *Ffar1* and *Ffar4* were broadly expressed in several enteroendocrine cell lineages, but largely excluded from enterochromaffin cells (Figure 2D). Orthogonally, the toxin receptor gene *Trpa1* was enriched in enterochromaffin cells (Bellono et al., 2017), but not abundantly expressed in other enteroendocrine cells (Figure 2D, E). Enterochromaffin cells also reportedly sense force through the mechanosensory ion channel PIEZO2 (Alcaino et al., 2018); *Piezo2* transcript was not readily detected in our transcriptomic data, but perhaps is enriched in enteroendocrine cells from colon which we did not analyze (Treichel et al., 2022) (Figure 2D). Thus, enteroendocrine cells often express multiple cell surface receptors, suggesting polymodal response properties, and some receptors are expressed by multiple enteroendocrine cell types.

### Genetic access to subtypes of enteroendocrine cells

Next, we obtained genetic tools for selective access to each major enteroendocrine cell lineage. We chose several combinations of Cre and FlpO lines to achieve intersectional genetic access to different enteroendocrine cells based on the cell atlas. (1) *Vil1-Cre*; *Pet1-FlpE* mice broadly target enterochromaffin cells, while (2) *Tac1-ires2-Cre; Vil1-p2a-FlpO* and (3) *Npy1r-Cre; Vil1-p2a-FlpO* mice target different enterochromaffin cell subtypes. (4) *Vil1-Cre; Sst-ires-FlpO*, (5) *Gip-Cre; Vil1-p2a-FlpO*, (6) *Cck-ires-Cre; Vil1-p2a-FlpO*, and (7) *Gcg-Cre; Vil1-p2a-FlpO* mice respectively target D, K, I, and L cells (Figure 2E).

Mice of each intersectional allele combination were crossed to *inter-tdTomato* mice, and reporter expression was analyzed across tissues, including in the brain, tongue, airways, pancreas, stomach, and intestine (duodenum to rectum) (Figure 3 – figure supplement 1, 2). Each of these seven intersectional combinations produced sparse labeling of intestinal epithelial cells, as expected for labeling of enteroendocrine cell subtypes (Figure 3 – figure supplement 1). Striking selectivity for enteroendocrine cells was observed across analyzed tissues for intersectional combinations targeting D, K, L, and I cells; sparse labeling was rarely observed in gastric endocrine cells and pancreatic islets, and absent from all other tissues examined. For example, *Cck-ires-Cre* alone (without intersectional genetics) drove reporter (*lsl-tdTomato*) expression in many tissues including the brain, spinal cord, and muscle, and within the intestine, in enteroendocrine cells as well as enteric neurons, extrinsic neurons, and cells in the lamina propria; however, in *Cck-ires-Cre; Vil1-p2a-FlpO*; *inter-tdTomato* mice, expression was not observed in the brain, spinal cord, or muscle, and within the intestine, was highly restricted to a subset of enteroendocrine cells, and not observed in other intestinal cell types (Figure 3 – figure supplement 1). We did note that *Tac1-ires2-Cre; Vil1-p2a-FlpO* and *Npy1r-Cre; Vil1-p2a-FlpO* more broadly labeled rectal epithelium, and *Npy1r-Cre; Vil1-p2a-FlpO* additionally labeled taste cells as well as rare cells in the airways and epiglottis (Figure 3 – figure supplement 2, 3). We also note that other genetic tools were inefficient at targeting enteroendocrine cells, including *Nts-ires-Cre* and *Mc4r-t2a-Cre* mice (Figure 3 – figure supplement 3A).

Hormone expression can be dynamic in individual enteroendocrine cells, and Cre/Flp lines provide an indelible marker for transiently expressed genes (Beumer *et al.*, 2018; Gehart *et al.*, 2019). Thus, Cre/Flp lines enable *in vivo* lineage tracing to measure enteroendocrine cell dynamics. Next, we used two-color expression analysis to investigate the repertoire of enteroendocrine cells captured by different Cre lines. Intestine cryosections were obtained from *Sst-ires-Cre*; *lsl-tdTomato*, *Pet1-Flpe; fsf-Gfp*, *Tac1-ires2-Cre*; *lsl-tdTomato*, *Npy1r-Cre*; *lsl-tdTomato*, and *Gcg-Cre*; *lsl-tdTomato*, *Cck-ires-Cre*; *lsl-tdTomato* mice (Figure 3, Figure 3 – figure supplement 3A). Two-color analysis involved visualization of native reporter fluorescence and immunohistochemistry for GLP1, CCK, SST, and/or serotonin in the duodenum. *Sst-ires-Cre* and *Pet-FlpE* mice showed enriched targeting of somatostatin and serotonin cells respectively (*Sst-ires-Cre* cells: 98.0% express somatostatin, 1.2% express serotonin, 0.8% express CCK, and 0.0% express GLP1; *Pet-FlpE* cells: 2.8% express somatostatin, 84.3% express serotonin, 19.1% express CCK, and 9.5% express GLP1) (Figure 3C). We note that the *Pet-FlpE* driver also captured a minority of cells with other hormones, suggesting that rare enteroendocrine cells can either transiently express markers of multiple lineages earlier in development or can switch identity from enterochromaffin cells to other enteroendocrine cell types. *Tac1-ires2-Cre* and *Npy1r-Cre* both labeled subsets of serotonin cells (100% of labeled cells produce serotonin in each line), with *Tac1-ires2-Cre* labeling a higher percentage of serotonin cells (78.4%) than *Npy1r-Cre* (5.0%) (Figure 3 – figure supplement 3). Both *Gcg-Cre* and *Cck-ires-Cre* mice labeled the majority of GLP1 and cholecystokinin cells; these cell types are within the same developmental lineage, and CCK and proglucagon are frequently coexpressed in the same EE cells (Habib et al., 2012). *Gcg-Cre* mice did not effectively label either somatostatin or serotonin cells (*Gcg-Cre* labeled 93.0% of GLP1 cells, 62.2% of CCK cells, 0.0% of somatostatin cells, and 0.6% of serotonin cells). *Cck-ires-Cre* mice were less selective (*Cck-ires-Cre* labeled 73.9% of GLP1 cells, 79.3% of CCK cells, 20.0% of somatostatin cells, and 6.0% of serotonin cells), and a substantial fraction (at least 23.8%) targeted other enteroendocrine cells that do not express these four hormones (Figure 3C). It is possible that the *Cck-ires-Cre* allele simply displays inefficient targeting efficiency, and/or that it drives reporter expression at early developmental time points with subsequent switching or refinement of cell identity. Together, these experiments measure the extent of selectivity achievable with each genetic tool, with some intersectional combinations providing highly selective genetic access to classes of enteroendocrine cells *in vivo*.

**Figure 3.**
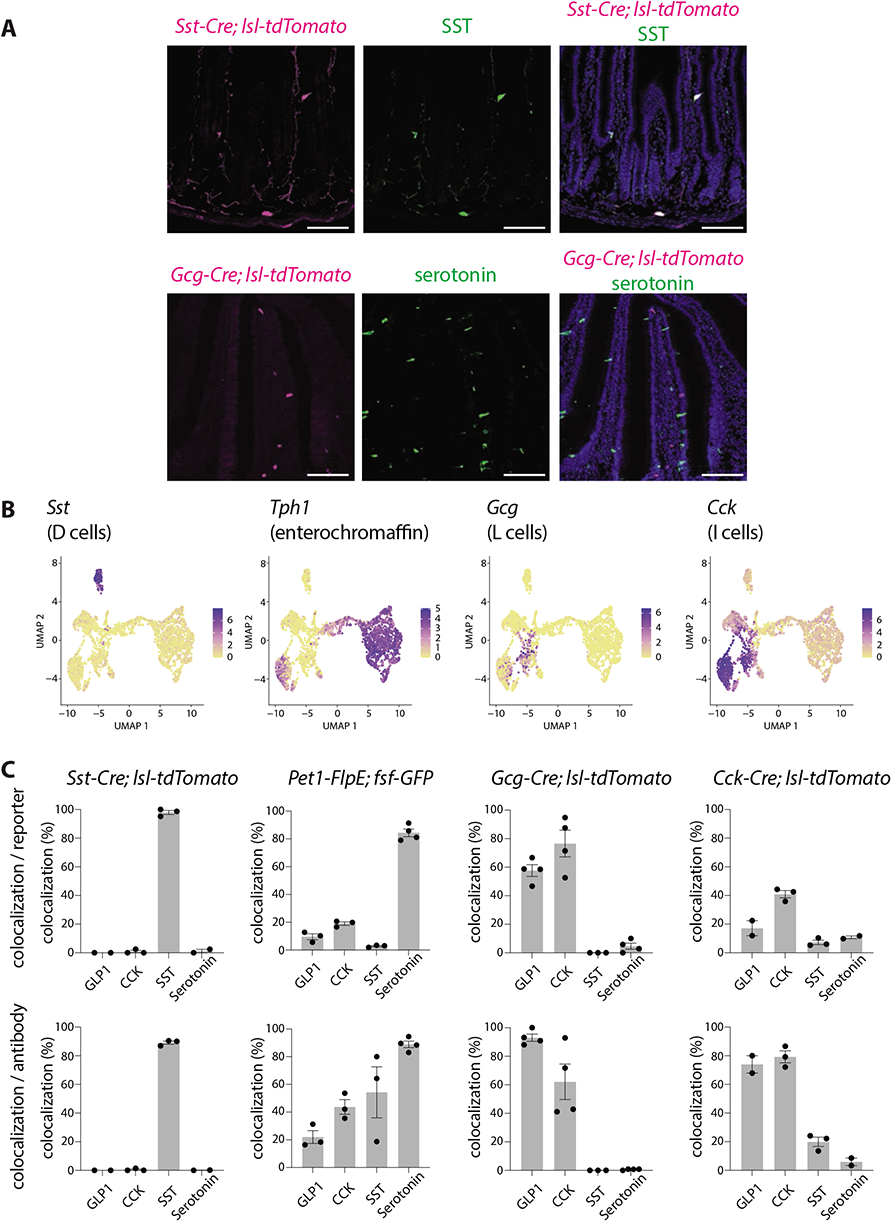
Differential targeting of enteroendocrine cell types by Cre lines. (A) Intestinal tissue was harvested from Cre lines indicated and analyzed simultaneously for expression of tdTomato (native fluorescence, red) and hormones (immunohistochemistry, green), scale bars: 100 μm. (B) Uniform manifold approximation and projection (UMAP) plots based on single-cell transcriptome data showing expression of indicated genes across the enteroendocrine cell atlas. (C) Quantification of cells expressing both Cre-based reporter and hormones indicated are shown as a percentage of all reporter-expressing cells (top) or all hormone-expressing cells (bottom), dots: individual animals, n: 2-4 mice, 14-901 cells/mouse (see Source Data for more information), mean ± sem. See Figure 3 - figure supplements 1, 2, and 3.

### Physiological responses to enteroendocrine cell activation

Direct study of enteroendocrine cell function has been challenging due to a lack of specific genetic tools. Neurogenin3 point mutations in human infants or intestine-targeted *Neurog3* knockout causes loss of enteroendocrine cells, severe malabsorptive diarrhea, and increased mortality (Mellitzer et al., 2010; Wang et al., 2006). We sought to develop cell type-specific genetic tools for enteroendocrine cell manipulation, reasoning that they might provide a complementary and specific approach to define the repertoire of evoked physiological and behavioral responses.

We first developed chemogenetic approaches for acute stimulation of all enteroendocrine cell types in freely behaving mice. Chemogenetic strategies involved designer G protein-coupled receptors (so-called DREADDs) that respond to the synthetic ligand clozapine-N-oxide (CNO) (Roth, 2016). *Neurod1-Cre*; *Vil1-p2a-FlpO* mice were crossed to contain an intersectional reporter allele (*inter-hM3Dq-mCherry*) that enables expression of a G*α*_q_-coupled DREADD (hM3Dq) only in cells expressing both Cre and Flp recombinase (Sciolino et al., 2016). Since this approach yielded rare reporter expression in pancreatic islets, we used an additional control mouse line, *Ptf1a-Cre*; *Vil1-p2a-FlpO*, which targets sparse *Vil1*-expressing pancreatic cells but not intestinal cells (Figure 4 – figure supplement 1) (Kawaguchi et al., 2002).

First, we examined the effect of global enteroendocrine cell activation on gut motility as assessed by movement of charcoal dye following oral gavage. *Neurod1-Cre*; *Vil1-p2a-FlpO*; *inter-hM3Dq-mCherry* mice, *Ptf1a-Cre; Vil1-p2a-FlpO*; *inter-hM3Dq-mCherry* mice, and control Cre-negative *Vil1-p2a-FlpO*; *inter-hM3Dq-mCherry* littermates were injected intraperitoneally (IP) with CNO (fed *ad libitum*, daytime). After 15 minutes, charcoal dye was administered, and after an additional 20 minutes, the gastrointestinal tract was harvested. Charcoal transit distance was calculated by genotype-blinded measurement of the charcoal dye leading edge. In control animals lacking DREADD expression, the leading edge of charcoal dye traversed part of the intestine (littermate controls lacking *Neurod1-Cre*: 22.6 ± 1.2 cm; littermate controls lacking *Ptf1a-Cre*: 22.8 ± 2.0 cm) (Figure 4). Chemogenetic activation of enteroendocrine cells accelerated gut transit, with the charcoal leading edge traversing 30.8 ± 1.5 cm of the intestine following CNO-injection in *Neurod1-Cre*; *Vil1-p2a-FlpO*; *inter-hM3Dq-mCherry* mice. When DREADD signaling was instead activated in all epithelial cells using *Vil1-Cre; lsl-hM3Dq* mice, gavaged dye failed to enter the intestine at all (Figure 4 – figure supplement 1A). CNO-accelerated gut transit was not observed in *Ptf1a-Cre; Vil1-p2a-FlpO*; *inter-hM3Dq-mCherry* mice (22.6 ± 2.6 cm) containing DREADD expression only in pancreatic cells (Figure 4, Figure 4– figure supplement 1B, C). Based on these observations, the observed effects in *Neurod1-Cre*; *Vil1-p2a-FlpO*; *inter-hM3Dq-mCherry* mice are due to enteroendocrine cells rather than pancreatic cells, and the net effect of activating all enteroendocrine cells is to promote gut transit.

**Figure 4.**
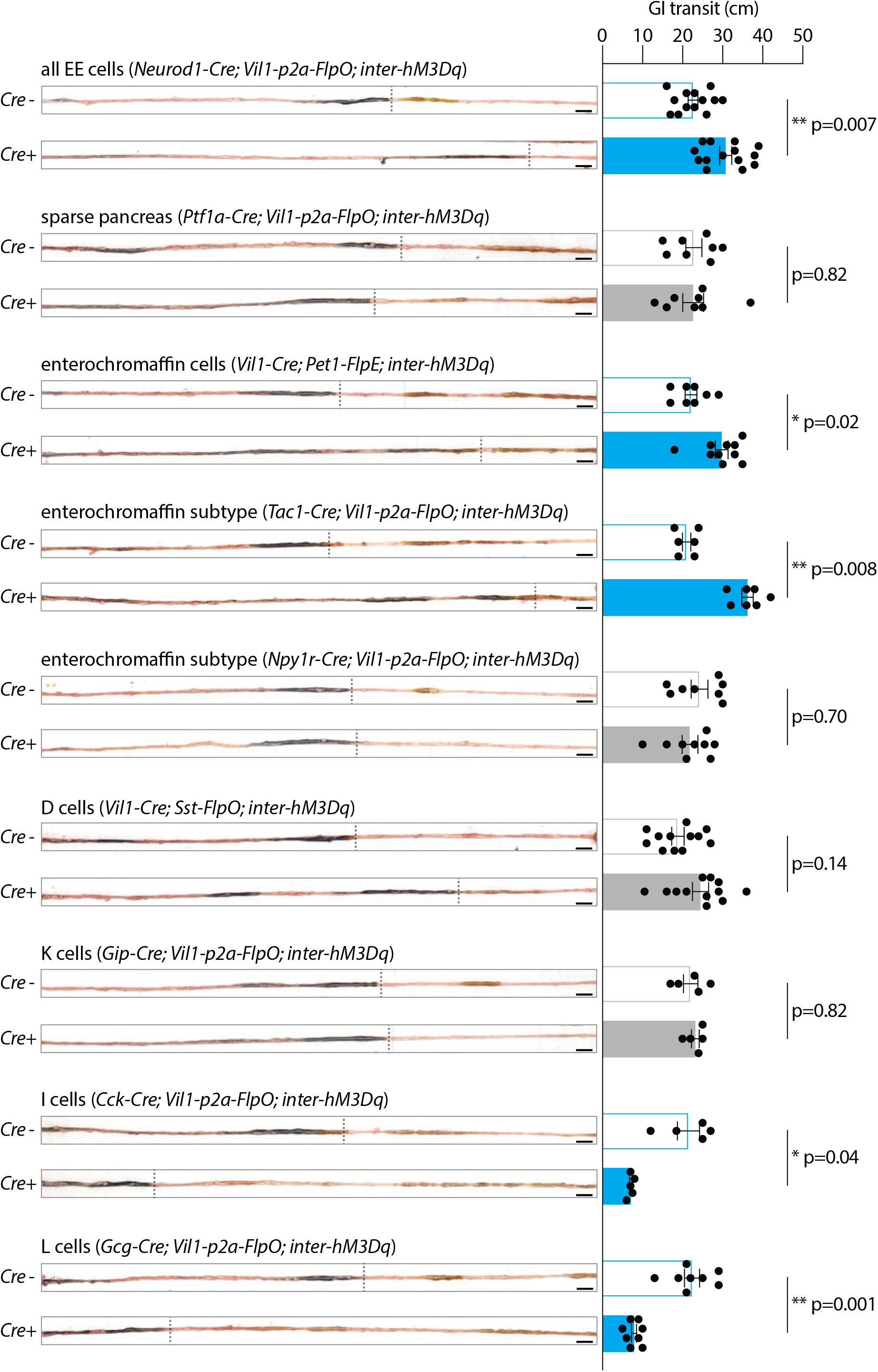
Enteroendocrine cell types that accelerate or slow gut transit. Mice of genotypes indicated were injected with CNO (IP, 3 mg/kg), and gavaged orally with charcoal dye. Intestinal tissue was harvested, and the distance between the pyloric sphincter and the charcoal dye leading edge was measured. Representative images (left) and quantification (right) of gut transit, scale bar: 1 cm, circles: individual mice, n: 5-14 mice, mean ± sem, *p<0.05, **p<0.01 by a Mann-Whitney test with Holm-Šídák correction. See Figure 4-figure supplement 1.

Next, we examined the effects of activating different enteroendocrine cell subtypes on gut motility. We additionally generated (1) *Vil1-Cre*; *Pet1-FlpE*; *inter-hM3Dq-mCherry* (2) *Tac1-ires2-Cre; Vil1-p2a-FlpO*; *inter-hM3Dq-mCherry*, (3) *Npy1r-Cre; Vil1-p2a-FlpO*; *inter-hM3Dq-mCherry*, (4) *Vil1-Cre; Sst-ires-FlpO*; *inter-hM3Dq-mCherry*, (5) *Gip-Cre; Vil1-p2a-FlpO*; *inter-hM3Dq-mCherry*, (6) *Cck-ires-Cre; Vil1-p2a-FlpO*; *inter-hM3Dq-mCherry*, and (7) *Gcg-Cre; Vil1-p2a-FlpO*; *inter-hM3Dq-mCherry* mice, with Cre-negative FlpO-positive *inter-hM3Dq-mCherry* littermates serving as controls (Figure 4). As above, CNO was injected (IP) into *ad libitum* fed animals followed by oral charcoal gavage. Enteroendocrine cells that produce serotonin promoted gut transit (*Pet-FlpE*: 29.8 ± 1.6 cm, Cre negative littermates: 22.1 ± 1.5 cm), while somatostatin-and GIP-expressing cells had no significant effect (*Sst-ires-FlpO*: 24.5 ± 2.0 cm, Cre negative littermates: 18.8 ± 1.6 cm; *Gip-Cre*: 23.2 ± 1.0 cm, *Gip-Cre* negative littermates: 22.0 ± 1.8 cm). Interestingly, single cell transcriptome data revealed multiple subtypes of enterochromaffin cells, and we observed accelerated gut transit upon chemogenetic activation of *Tac1*-expressing cells (*Tac1-ires2-Cre*: 36.2 ± 1.4 cm, *Tac1-ires2-Cre* negative littermates: 21.0 ± 1.1 cm) but not *Npy1r*-expressing cells (*Npy1r-Cre*: 21.8 ± 2.0 cm, *Npy1r-Cre* negative littermates: 24.3 ± 2.1 cm). These findings raise the possibility that each enterochromaffin cell subtype may privately communicate with different downstream extrinsic and/or enteric neurons to control gut physiology. In contrast, CCK-and GLP1-expressing cells slowed gut motility (*Cck-ires-Cre*: 7.1 ± 0.3 cm, *Cck-ires-Cre* negative littermates: 21.5 ± 2.8 cm; *Gcg-Cre*: 7.9 ± 0.7 cm, *Gcg-Cre* negative littermates: 22.4 ± 1.9 cm). Ingested food slows gut motility to promote nutrient absorption, while ingested toxins may accelerate gut motility to purge luminal contents (Nozawa et al., 2009; Van Citters and Lin, 2006). Consistent with these findings, CCK and GLP1 are released by nutrients while serotonin signaling is required for certain toxin responses (Drucker, 2016; Gribble and Reimann, 2019). Simultaneous activation of both pathways, as done in *Neurod1-Cre*; *Vil1-p2a-FlpO*; *inter-hM3Dq-mCherry* mice, masks the slowing of gut transit by CCK and GLP1 cells. These findings suggest a hierarchy where neural circuits that mediate toxin responses may achieve priority over those that mediate nutrient responses, at least under conditions of equal and maximal activation. Altogether, we characterize enteroendocrine cell subtypes that have different and sometimes opposing effects on digestive system physiology.

### Enteroendocrine cells that regulate feeding and nausea-associated behavior

Next, we examined the effect of global enteroendocrine cell activation on feeding behavior. Fasted *Neurod1-Cre*; *Vil1-p2a-FlpO*; *inter-hM3Dq-mCherry* mice, *Ptf1a-Cre; Vil1-p2a-FlpO*; *inter-hM3Dq-mCherry* mice, and control littermate mice lacking Cre recombinase were injected (IP) with CNO and given access to food for two hours at dark onset (Figure 5A). Animals lacking DREADD expression, or with DREADD expression only in pancreas, ate robustly (∼1 g of food over a two-hour period). In contrast, CNO-induced activation of enteroendocrine cells in *Neurod1-Cre*; *Vil1-p2a-FlpO*; *inter-hM3Dq-mCherry* mice caused a 26% reduction in food intake (Figure 5B).

**Figure 5.**
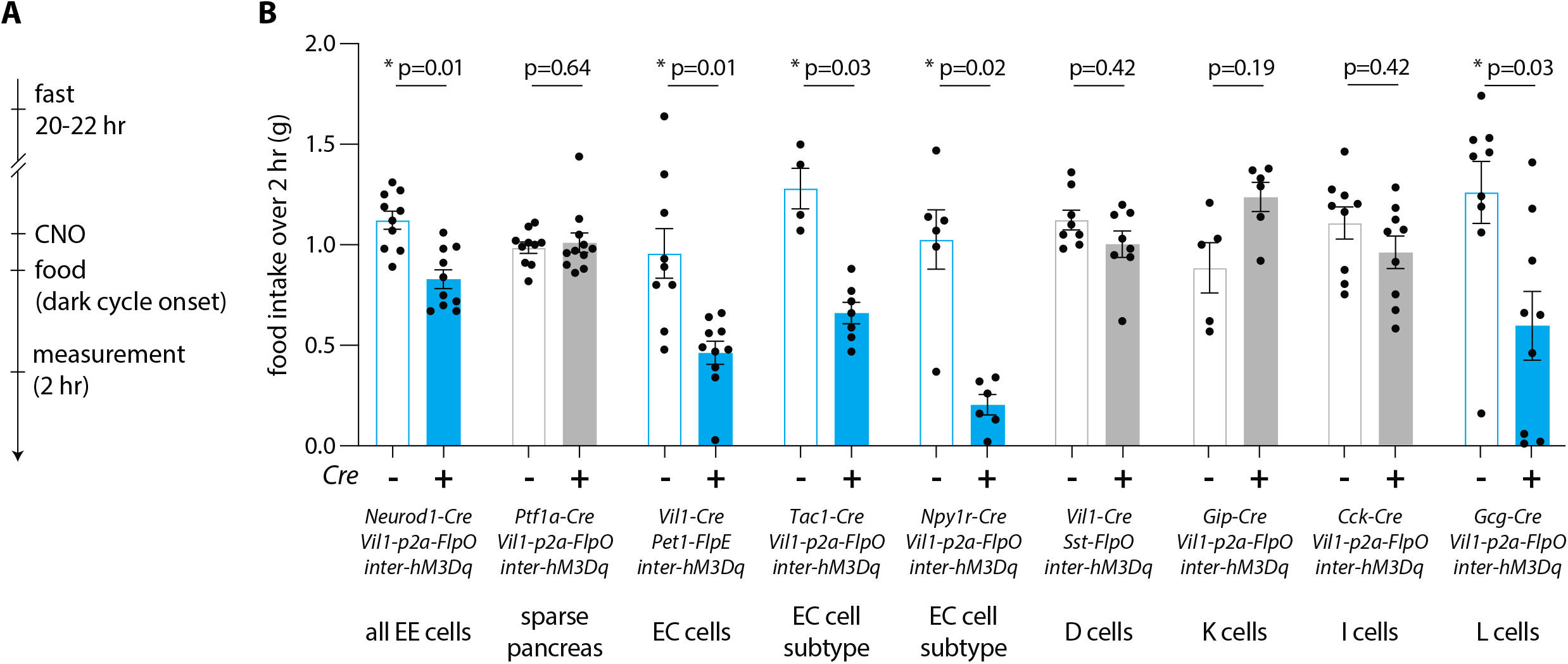
Enteroendocrine cell types that reduce feeding. (A) Timeline for behavioral assay. (B) Mice of genotypes indicated were fasted overnight, injected with CNO (IP, 3 mg/kg), and total food intake was measured during two hour *ad libitum* food access, circles: individual mice, n: 6-11 mice, mean ± sem, *p<0.05 by a Mann-Whitney test with Holm-Šídák correction. See Figure 5 - figure supplement 1.

To interrogate the roles of different enteroendocrine cell subtypes in feeding regulation, similar experiments were then performed in (1) *Vil1-Cre*; *Pet1-FlpE*; *inter-hM3Dq-mCherry* (2) *Tac1-ires2-Cre; Vil1-p2a-FlpO*; *inter-hM3Dq-mCherry*, (3) *Npy1r-Cre; Vil1-p2a-FlpO*; *inter-hM3Dq-mCherry*, (4) *Vil1-Cre; Sst-ires-FlpO*; *inter-hM3Dq-mCherry,* (5) *Gip-Cre; Vil1-p2a-FlpO*; *inter-hM3Dq-mCherry*, (6) *Cck-ires-Cre; Vil1-p2a-FlpO*; *inter-hM3Dq-mCherry*, and (7) *Gcg-Cre; Vil1-p2a-FlpO*; *inter-hM3Dq-mCherry* mice, with Cre-negative, FlpO-positive, *inter-hM3Dq-mCherry* littermates again serving as controls. Chemogenetic activation of all *Pet1*-expressing enterochromaffin cells, and enterochromaffin cell subtypes marked in *Tac1-ires2-Cre* and *Npy1r-Cre* mice, reduced feeding behavior (compared to Cre-negative littermates, *Pet1-FlpO*: 52.1% reduction, *Tac1-ires2-Cre*: 48.5% reduction, *Npy1r-Cre*: 79.6% reduction), while activation of D and K cells did not (Figure 5B). Activating GLP1-producing L cells also reduced feeding (compared to Cre-negative littermates, *Gcg-Cre*: 52.4% reduction), but surprisingly, activating CCK-producing I cells lowered feeding only in fed but not fasted mice (Figure 5 – figure supplement 1A). This observation is likely due to *Cck-ires-Cre* and *Gcg-Cre* alleles targeting at least partially distinct populations of enteroendocrine cells. Chemogenetic activation of L cells (single CNO injection) caused a durable reduction of feeding for several hours, with total food intake normalizing by 11 hours, and also evoked a decrease in water intake but not locomotion (Figure 5 – figure supplement 1B). For comparison, activating somatostatin cells did not change feeding, water intake, or locomotion. Altogether, we find that some but not all enteroendocrine cells can regulate food intake, and can do so with varying efficacy.

Feeding behavior can be suppressed by a variety of stimuli, including nutrients that induce satiety (the feeling of fullness) and toxins that induce the unpleasant sensation of nausea. Satiety and nausea can be distinguished in mice using classical learning paradigms, as nutrients provide positive reinforcement signals while nausea-inducing toxins cause learned avoidance of paired flavors (de Araujo et al., 2008; Fernandes et al., 2020; Han et al., 2018; Prescott and Liberles, 2022; Tan *et al.*, 2020; Zhang et al., 2021). Here, we asked whether activating various enteroendocrine cells resulted in positive or negative reinforcement of behavioral responses to a novel flavor.

Conditioned flavor preference assays were performed to target enteroendocrine cell types that impact feeding behavior, along with somatostatin as a control. We obtained (1) *Vil1-Cre*; *Pet1-FlpE*; *inter-hM3Dq-mCherry* (2) *Vil1-Cre; Sst-ires-FlpO*; *inter-hM3Dq-mCherry,* (3) *Cck-ires-Cre; Vil1-p2a-FlpO*; *inter-hM3Dq-mCherry*, and (4) *Gcg-Cre; Vil1-p2a-FlpO*; *inter-hM3Dq-mCherry* mice, and *Cre*-negative littermates. Assays involved mice on a low-water regimen for three days, which ensured participation in behavioral assays. On a conditioning day, mice were given access to a drink with a novel flavor (either cherry or grape-flavored saccharin solution), and then immediately injected (IP) with CNO, saline, or the gut poison lithium chloride. On a subsequent testing day, mice were presented simultaneously with cherry and grape-flavored drink in a two-choice assay, and a preference index calculated as previously described (Figure 6A) (Zhang *et al.*, 2021). In the absence of malaise induction, control animals typically displayed a mild preference for the experienced flavor. In contrast, mice developed a strong aversion to flavors paired with lithium chloride injection (Figure 6 – figure supplement 1). Likewise, chemogenetic activation of GLP1-producing L cells provided a flavor avoidance teaching signal, and a trend was observed upon activating enterochromaffin cells (Figure 6B). In contrast, activation of D or I cells was without effect in this paradigm. These findings are consistent with prior observations that GLP1R agonists evoke nausea, while serotonin receptor blockade suppresses nausea responses to various poisons (Baggio and Drucker, 2007; Gale, 1995; Gribble and Reimann, 2021; Tyers and Freeman, 1992). Furthermore, we observed that flavor avoidance was promoted by enteroendocrine cell types with differential impact on gut motility, as TAC1 cells promoted gut transit, GLP1 cells decreased gut transit, and NPY1R cells had no effect. As with the feeding assay above, we observed different responses to activation of CCK and GLP1 cells, as *Cck-ires-Cre* and *Gcg-Cre* mice target only partially overlapping assortments of enteroendocrine cells. We note that enteroendocrine cells that mediate positive-reinforcement signals were not identified; perhaps this flavor conditioning paradigm is not sensitive enough to detect positive reinforcement signals, or perhaps other pathways are relevant for nutrient reward. Together, these findings indicate that particular enteroendocrine cells can inhibit feeding and also evoke nausea-associated behavior.

**Figure 6.**
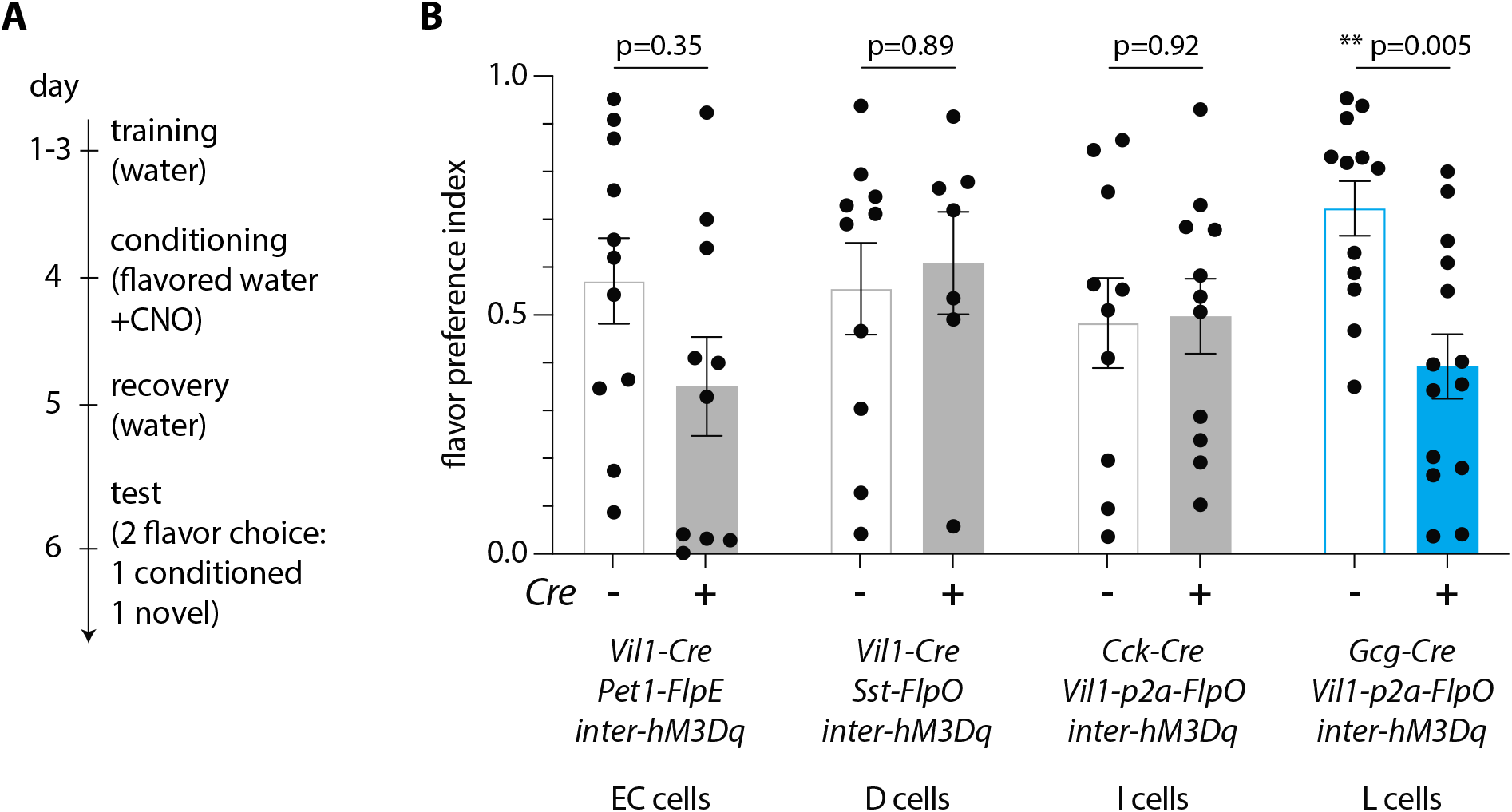
Enteroendocrine cell types that condition flavor avoidance. (A) Timeline for behavioral assay. (B) Mice of genotypes indicated were analyzed for CNO-evoked flavor avoidance, n: 7-14, mean ± sem, circles: individual data points, **p<0.01 by a Mann-Whitney test with Holm-Šídák correction. See Figure 6-figure supplement 1.

## CONCLUSION

Here we developed a toolkit involving intersectional genetics for systematic access to each major enteroendocrine cell lineage (Figure 7A). We then used chemogenetic approaches to delineate major response pathways of the gut-brain axis (Figure 7B). Serotonin-producing enterochromaffin cells express the irritant receptor TRPA1 (Bellono *et al.*, 2017) and chemogenetic activation blocks feeding behavior and promotes gut transit, presumably for toxin clearance. Furthermore, different enterochromaffin cell subtypes can have different effects on gut motility, suggesting at least partially nonoverlapping communication pathways with downstream neurons. These findings are consistent with a role for enterochromaffin cells in toxin-induced illness responses, and interestingly, pharmacological blockade of the serotonin receptor HTR3A is a clinical mainstay for nausea treatment (Freeman et al., 1992). Other enteroendocrine cell types, including those that produce CCK, GIP, GLP1, neurotensin, and somatostatin, express nutrient receptors yet elicit different physiological and behavioral responses. For example, GLP1 cells slow gut motility, presumably to promote nutrient absorption, decrease feeding behavior, and evoke conditioned flavor avoidance (Gribble and Reimann, 2019; Zhang *et al.*, 2021). Additional studies are needed to define gut-brain pathways that mediate nutrient reward, and why receptors for specific nutrients are expressed across a dispersed ensemble of enteroendocrine cells. Together, these experiments provide a highly selective method for accessing enteroendocrine cells in vivo, and a direct measure of their various roles in behavior and digestive physiology.

**Figure 7.**
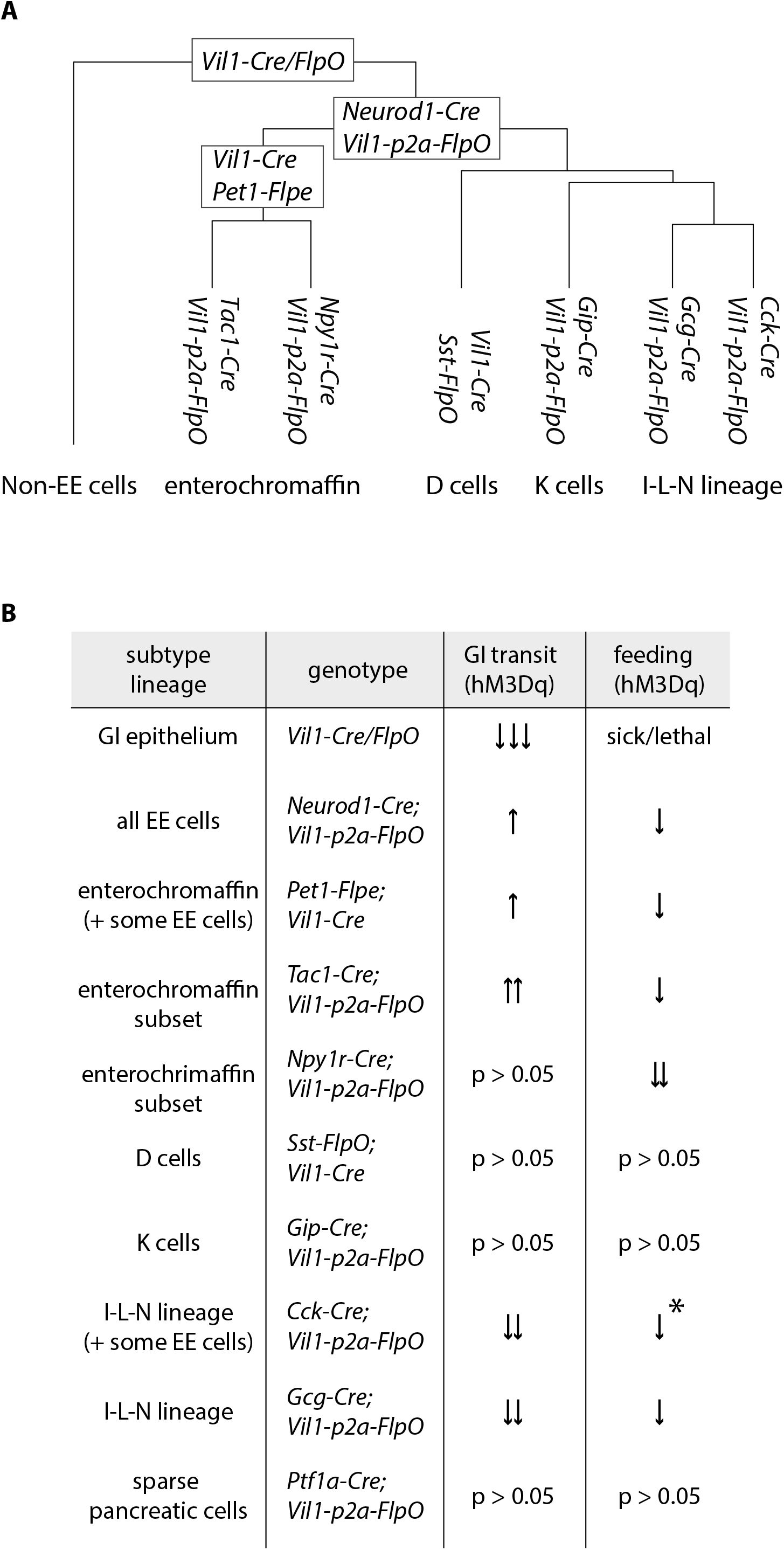
Differential regulation of physiology and behavior by enteroendocrine cell subtypes. (A) A dendrogram depicting cell types targeted by different genetic tools. (B) Summary of feeding and gut transit data obtained for genetic tools that target different enteroendocrine cell types, *only observed in fed state.

## ACKNOWLEDGMENTS

We thank Nancy Thornberry and Paul Richards for manuscript comments, Lin Gan (Rochester) for *Atoh1-Cre* mice, Chris Wright (Vanderbilt) for *Ptf1a-Cre* mice, Patricia Jensen (NIH) and Nicholas Plummer (NIH) for genotyping advice, members of the Liberles laboratory for experimental advice and assistance, the Boston Children’s Hospital Mouse Gene Manipulation Core (NIH P50 HD105351) for help with generating mice, the Beth Israel Deaconess Medical Center Energy Balance Core (OD028635, P30 DK034854) for help with CLAMS assays, the Harvard Medical School Nikon Imaging Center and Neurobiology Imaging Facility (NINDS P30 NS072030) for help with microscopy, the Biopolymers Facility for help with sequencing, the Immunology Flow Cytometry Core Facility for help with cell sorting, and the Harvard Medical School O2 High Performance Computer Cluster for bioinformatics support. This work was supported by the NIH (DP1AT009497 to SDL, R01DK103703 to SDL, and T32 HL007901 to JAK), and the Food Allergy Science Initiative (to SDL). MH is a fellow of Japan Society for the Promotion for Science. SDL is an investigator of the Howard Hughes Medical Institute.

## AUTHOR CONTRIBUTIONS

MH and SDL designed the study, analyzed data, and wrote the manuscript; MH performed all experiments, except ERD performed flavor avoidance assays involving chemogenetics and some two-color expression analyses and NRJ performed tongue and airway histology; JAK and MH analyzed transcriptome data; FR generated *Gip-Cre* mice.

## MATERIALS AND METHODS

### Mice

All animal husbandry and procedures were performed in compliance with institutional animal care and use committee guidelines. All animal husbandry and procedures followed the ethical guidelines outlined in the NIH Guide for the Care and Use of Laboratory Animals (https://grants.nih.gov/grants/olaw/guide-for-the-care-and-use-of-laboratory-animals.pdf), and all protocols were approved by the institutional animal care and use committee (IACUC) at Harvard Medical School. *Atoh1-Cre* knock-in (Yang et al., 2010), *Pet1-FlpE* (Jensen et al., 2008), and *Ptf1a-Cre* (Kawaguchi et al., 2002) mice were described before; *Atoh1-Cre* transgenic (011104), *Neurog3-Cre* (006333), *Neurod1-Cre* (028364), *Sst-ires-Cre* (013044), *Sst-ires-FlpO* (028579), *Vil1-Cre* (021504), *Gcg-Cre* (030542), *Cck-ires-Cre* (012706), *Nts-ires-Cre* (017525), *Mc4r-t2a-Cre* (030759), *Npy1r-Cre* (030544), *Tac1-ires2-Cre* (021877), *lsl-tdTomato* (*Ai14,* 007914), *inter-tdTomato* (*Ai65,* 021875), *inter-hM3Dq-mCherry* (*CAG-lsl-fsf-hM3Dq*, 026943), *lsl-hM3Dq* (026220), and C57BL/6 (000664) mice were purchased (Jackson Laboratory). Both male and female mice between 8-24 weeks old were used for all studies, and no differences based on sex were observed. All mice were maintained in the C57BL/6 genetic background. Mouse breeding involved paternal *Cre* alleles, paternal *Flp* alleles, and/or maternal effector genes. *Vil1-Cre* produced occasional germline recombination of *loxP* sites which resulted in ectopic *inter-hM3Dq-mCherry* gene expression; mice with such ectopic expression were excluded based on genotyping of reporter allele DNA extracted from ear tissue with primer 1 (stop cassette forward): atgtctggatctgacatggtaa; primer 2 (*hM3Dq* cassette reverse): tctggagaggagaaattgcca; primer 3 (GFP cassette reverse): ttgaagtcgatgcccttcag; intact allele: ∼490 bp, recombined allele: ∼290 bp. *Vil1-p2a-FlpO* mice were generated by CRISPR-guided approaches at Boston Children’s Hospital Mouse Gene Manipulation Core. Cas9 protein, CRISPR sgRNAs (targeting the stop codon of *Vil1* locus), and a ssDNA (containing a *p2a-FlpO* cassette with 150bp homology arms) were injected into the pronucleus of C57BL/6 embryos. Founder mice were screened by allele specific PCR analysis with primers flanking the 5’ junction (primer 1: aacagaagttccttaaacaagcca; primer 2: aacaggaactggtacagggtcttg; ∼930 bp), FlpO internally (primer 1: acaagggcaacagccaca; primer 2: tcagatccgcctgttgatgt; ∼830bp), and the 3’ junction (primer 1: accccctggtgtacctgga; primer 2: tagccctcccttttgagtgtga; ∼840 bp), followed by Sanger sequencing to validate the allele. Selected *Vil1-p2a-FlpO* founder mice were viable, fertile, and back crossed to C57BL/6 mice for at least 3 generations.

### Single-cell RNA sequencing

Enteroendocrine cells were acutely harvested using a protocol modified from previous publications (Haber *et al.*, 2017; Sato et al., 2009). Intestinal tissue was obtained from *Neurog3-Cre; lsl-tdTomato* mice (1 adult male), or *Neurod1-Cre; lsl-tdTomato* (3 adult females), cut longitudinally, washed (cold phosphate-buffered saline or PBS), cut into small ∼5 mm pieces, and incubated (gentle agitation, 20 minutes, 4°C) in EDTA solution (20 mM EDTA-PBS, Ca/Mg-free) in LoBind Protein tubes (Eppendorf 0030122216). The specimen was shaken, the tissue allowed to settle, and the supernatants collected. The residual tissue was again incubated similarly with EDTA solution, and supernatants were combined, and centrifuged (300 g, 5 minutes, 4°C) Pellets were washed (2x, PBS (Ca/Mg-free) supplemented with 5% Fetal Bovine Serum (FBS), 4°C) and incubated (37°C, 2 minutes) in protease solution (TrypLE express, Thermo Fisher 12604013) supplemented with DNase (100 U/ml, Worthington Biochemical LK003172). The suspension containing dissociated cells was centrifuged (300 g, 5 minutes), washed (2x, PBS (Ca/Mg-free) containing 5% FBS, 4°C) The resulting pellet was resuspended in FACS buffer (5% FBS in DMEM/F12, HEPES, no phenol red) containing DNase (100 U/ml), TO-PRO-3 (Thermo Fisher T3605, 1:10,000) to label dead cells, and Calcein Violet (Thermo Fisher 65-0854-39, 1:10,000) to label living cells. Cells were filtered (1 x 70-um, 1 x 40-um) and tdTomato+, Calcein Violet+, TO-PRO-3-cells were collected by fluorescence activated cell sorting using a FACS Aria (BD Biosciences). Collected cells were then loaded into the 10x Genomics Chromium Controller, and cDNA prepared and amplified according to manufacturer’s protocol (10x Genomics, Chromium single cell 3’ reagent kit v3, 12 cycles per amplification step). The resulting cDNA was sequenced on a NextSeq 500 at the Harvard Medical School Biopolymers Facility. Sequence reads were aligned to the mm39 mouse transcriptome reference, and feature barcode matrices were generated using 10x Genomics CellRanger. Unique transcript (UMI) count matrices were analyzed in R v4.1.1 using Seurat v4.0.5 (Beutler et al., 2017; Satija et al., 2015). The cell barcodes were filtered, removing cells with a high number of UMIs (>125,000) or high percentage of mitochondrial genes (>25%). The filtered UMI count matrix was transformed using SCTransform (Hafemeister and Satija, 2019). Transformed matrices from *Neurog3* and *Neurod1* samples were integrated (nFeature = 3000), and integrated matrices used for cluster identification and UMAP projections. Additional clusters of low-quality cells (defined by low average UMI counts and low average feature counts across the cluster) were removed. To examine the diversity among enteroendocrine cells, cell barcodes belonging to enteroendocrine cells from *Neurog3* and *Neurod1* samples were identified and reanalyzed separately. Matrices of enteroendocrine cells from *Neurog3* and *Neurod1* samples were transformed and integrated (nFeature = 3000). Differential gene expression (Wilcoxon-Ranked Sum test) was conducted on UMI counts matrices that were log normalized and scaled. Seurat’s BuildClusterTree function was used to spatially arrange clusters based on relative similarity in gene expression. Two serotonergic clusters were merged *post hoc* (to become cluster EC_3) due to the absence of any single signature gene that effectively distinguished them. Gene expression data in all UMAP plots is shown as a natural log of normalized UMI counts. Further details and full parameters of analysis will be provided on GitHub upon publication: https://github.com/jakaye/EEC_scRNA.

### Tissue histology

For histology, mice were perfused intracardially with PBS and then fixative (4% paraformaldehyde/PBS), and intestinal regions and other organs dissected (duodenum: first 2 cm after the pyloric sphincter, jejunum: middle 2 cm, and ileum: last 2 cm before the cecum) and postfixed (1-2 hours, 4°C). Samples were then incubated in 30% sucrose/PBS (overnight, 4°C), embedded in Tissue-Tek OCT, frozen, cryosectioned, and placed on glass slides. Slides were incubated with primary antibodies at dilutions indicated below (overnight, 4°C, PBS supplemented with 0.05% Tween20, 0.1% TritonX, and either 5% normal donkey serum or 1% BSA) and then with fluorophore-conjugated secondary antibodies (1:500, 2 hours, RT). Sections were mounted (DAPI Fluoromount-G, Southern Biotech 0100-20), coverslipped, and imaged using a Nikon A1R confocal microscope, an Olympus FV1000 confocal microscope, or a Zeiss Axiozoom.V16 fluorescent stereoscope. Microscope images are presented as z-projections. Two-color images (Nikon Ti2 inverted microscope) were analyzed using the cell counter function on the Nikon NIS-element software. Antibodies were: Rabbit anti-CCK (Abcam ab27441, 1:1000), Rabbit anti-CRE (Cell Signaling 15036, 1:500), Rabbit anti-GLP1 (Novus 2622B MAB10473, 1:2000), Rabbit anti-NTS (Immunostar 20072, 1:2000), Rabbit anti-SST (Novus 906552 MAB2358, 1:1000), Goat anti-5HT (Abcam ab66047, 1:2000), Donkey anti-rabbit Alexa488, Cy3, Cy5, Alexa680 (Jackson Immuno Research, ThermoFisher, 1:500), Donkey anti-goat Alexa488 (Jackson Immuno Research, 1:500).

### Gut transit measurements

DREADD-expressing and control animals (*ad libitum* fed) were injected with CNO (3 mg/kg, IP). After 15 minutes, charcoal dye (200 μl, 10% activated charcoal, 10% gum Arabic in water), or for Figure 4 – figure supplement 1, carmen red dye (200 μl, 6% carmen red, 0.5% methyl cellulose in water), was gavaged orally, and 20 minutes later, mice were euthanized and the gastrointestinal tract was harvested. The distance between the pyloric sphincter and the charcoal dye leading edge was measured by an observer blind to animal genotype. All animals were naive to CNO exposure, except for some *Gip-Cre* mice due to limited availability of mice.

### Feeding measurements

Experimental mice were individually housed for 3 days, and habituated to feeding from a ceramic bowl. Animals were either fed *ad libitum* or fasted for the last 20-22 hours in a new clean cage with some bedding material from the previous cage. CNO was injected (3 mg/kg, IP), and food pellets presented 15 minutes later at the onset of darkness. Food intake was measured over the course of 2 hours by weighing the amount of residual food, with genotypes revealed *post hoc* to achieve a genotype-blinded analysis. Studies involved fasted mice that were naive to prior CNO exposure or fed mice that were either naive to CNO or acclimated for at least a week after prior CNO exposure.

### Body Composition and Indirect Calorimetry

Body composition (lean mass and fat mass) was first analyzed for each experimental group with a 3-in-1 Echo MRI Composition Analyzer (Echo Medical Systems, Houston, Texas), and no significant differences were observed. Animals were then placed in a Sable Systems Promethion indirect calorimeter maintained at 23°C ± 0.2°C. Mice were singly housed in metabolic cages with corn cob bedding and *ad libitum* access to Labdiet 5008 chow (56.8/16.5/26.6 carbohydrate/fat/protein). After 18 hours, all mice were injected with PBS (IP) for acclimatation to handling and mild injection stress. The following day, mice were injected with CNO (3 mg/kg, IP) approximately 15 minutes before dark onset. Animals were then analyzed for food and water consumption, body weight, and distance traveled. Statistical analysis was performed with CalR (Mina et al., 2018).

### Conditioned flavor avoidance

Conditioned flavor avoidance assays were performed over a 6-day protocol as described previously (Zhang *et al.*, 2021). Mice were water restricted in their home cage throughout the assay, and given brief *ad libitum* water access (30 minutes) only in testing arenas to ensure participation in behavioral trials. On days 1-3, mice were housed individually and each day were introduced to a test arena (30 minutes, dark onset) where they were given brief *ad libitum* access to water from two water bottles. On day 4 (conditioning day), mice were introduced (30 minutes) to a test arena containing 2 bottles of either grape-flavored or cherry-flavored Kool-Aid supplemented with 0.2% saccharin for 30 minutes, and then immediately injected (IP) with CNO (3 mg/kg), saline, or LiCl (0.6 M, saline) and returned to the home cage. On day 5 (recovery day), mice (like days 1-3) were introduced into a test arena (30 minutes, dark onset) where they were given brief *ad libitum* access to water from two water bottles. On day 6 (testing day), mice were given one bottle of cherry-flavored water with 0.2% saccharin and one bottle of grape-flavored water with 0.2% saccharin on randomized sides, and consumption from each bottle was measured using an automated lickometer for 30 minutes (Slotnick, 2009; Zhang *et al.*, 2021). Analysis of LiCl responses involved a house-made automated lickometer using commercial components (Arduino Uno A000066, Adafruit MPR121 Capacitive Sensor 1982). Genotypes were revealed *post hoc* to avoid potential bias.

### Statistical analysis

Graphs represent data as mean ± SEM, as indicated in figure legends. All data points were derived from different mice except some mice in Figure 4 (*Gip-Cre*; *Vil1-p2a-FlpO*; *inter-hM3Dq-mCherry* mice) were previously used in feeding assays and some mice in Figure 5-figure supplement 1 (*Ptf1a-Cre; Vil1-p2a-FlpO; inter-hM3Dq-mCherry*: 21/21 mice, *Cck-ires-Cre; Vil1-p2a-FlpO; inter-hM3Dq-mCherry*: 7/19 mice, *Gip-Cre; Vil1-p2a-FlpO; inter-hM3Dq-mCherry*: 10/10 mice, and *Vil1-Cre; Sst-ires-FlpO; inter-hM3Dq-mCherry*: 9/16 mice) were previously used in prior feeding assays for Figure 5. When mice were reused, they were acclimated for at least a week after prior CNO exposure.

Sample sizes (from left to right): Figure 3C (Sst: 2, 3, 3, 2; Pet1: 3, 3, 3, 4; Gcg: 4, 4, 3, 4; Cck: 2, 3, 3, 2), Figure 4 (13, 14, 8, 8, 8, 10, 6, 7, 8, 9, 12, 12, 5, 5, 5, 5, 8, 8), Figure 4 – figure supplement A (4, 5), Figure 5 (10, 10, 10, 11, 9, 10, 4, 7, 6, 6, 8, 8, 5, 6, 9, 9, 9, 9), Figure 5 –figure supplement 1A (9, 12, 9, 8, 4, 6, 9, 10), Figure 5 – figure supplement 1B (8, 8, 4, 4), Figure 6 (11, 10, 10, 7, 10, 11, 12, 14), Figure 6 – figure supplement 1 (11, 11).

Statistical significance was measured using a Mann-Whitney test with Holm-Šídák correction on Prism 9 (Graphpad) for Figure 4, Figure 5, Figure 5 – figure supplement 1A, and Figure 6, a Mann-Whitney test on Prism 9 (Graphpad) for Figure 4-figure supplement 1A and Figure 6-figure supplement 1A, and ANCOVA and ANOVA on CalR for Figure 5-figure supplement 1B (Mina *et al.*, 2018).

### Source data

The source data excel file contains raw numerical data used for all bar graphs and statistical analysis. Single-cell transcriptome data will be available with a GEO GSE accession number upon acceptance of the manuscript.

### Material availability statement

*Vil1-p2a-FlpO* mice will be deposited in Jackson Laboratory and made generally available upon reasonable request.

---

**Figure 1 - figure supplement 1.**
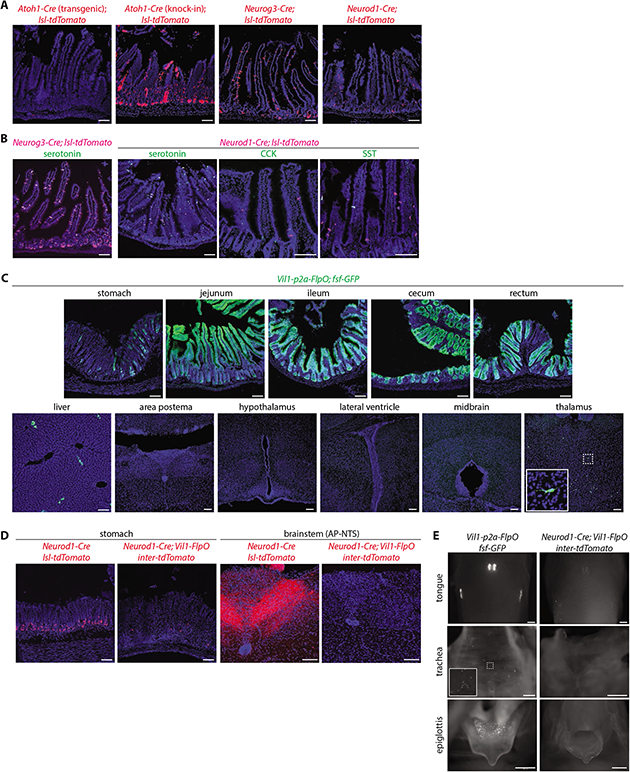
Characterization of mouse lines for intersectional genetics. (A) Native tdTomato fluorescence in fixed intestinal cryosections (20 μm) of mouse lines indicated. (B) Two-color analysis examining expression of tdTomato (native fluorescence, magenta) and hormones (immunochemistry, green) in fixed intestinal cryosections (20 μm) of mouse lines indicated, SST: somatostatin. (C) Native GFP fluorescence in fixed cryosections of tissues (20 μm) and brain regions indicated (50 μm) in *Vil1-p2a-FlpO; fsf-Gfp* mice. (D) Native tdTomato fluorescence in fixed tissue cryosections (20 μm) from mouse lines indicated. (E) Native reporter fluorescence in fixed wholemount tissue preparations from mouse lines indicated. Scale bars: 100 μm.

**Figure 2 - figure supplement 1.**
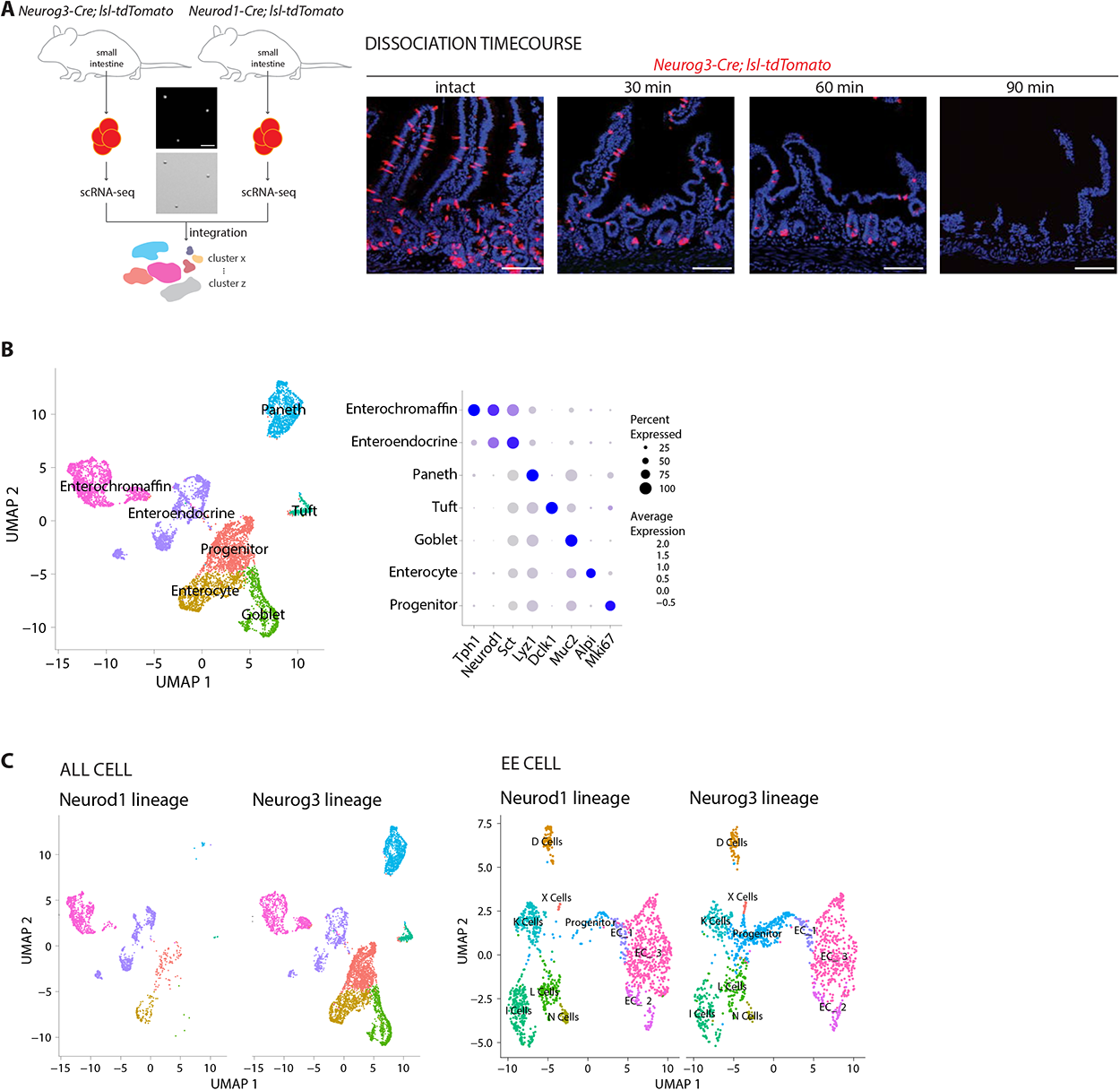
Analyzing cell types marked in different Cre-defined lineages. (A) Left: small intestines were collected from *Neurog3-Cre; lsl-tdTomato* (1 male) and *Neurod1-Cre; lsl-tdTomato* (3 female) mice. Fluorescent cells were collected by cell sorting and separately analyzed by single-cell RNA sequencing. (Center) Cell purity was determined by visualizing tdTomato fluorescence (top) and bright field microscopy (bottom), scale bar: 100um. Right: duodenum tissue from *Neurog3-Cre; lsl-tdTomato* mice was fixed at different time points after the onset of cell dissociation, and analyzed for tdTomato fluorescence in 20 μm tissue cryosections, scale bars: 100 μm. (B) (left) A uniform manifold approximation and projection (UMAP) plot of single-cell transcriptomic data merged from both *Neurog3-Cre; lsl-tdTomato* and *Neurod1-Cre; lsl-tdTomato* mice. Cells are colored based on the expression of signature genes indicated in the dot plot on right (average expression from natural log of normalized UMI counts). (C) UMAP plots indicating all cell types (left) and enteroendocrine cell types (right) purified from *Neurod1-Cre; lsl-tdTomato* (Neurod1 lineage) and *Neurog3-Cre; lsl-tdTomato* (Neurog3 lineage) mice.

**Figure 3 - figure supplement 1.**
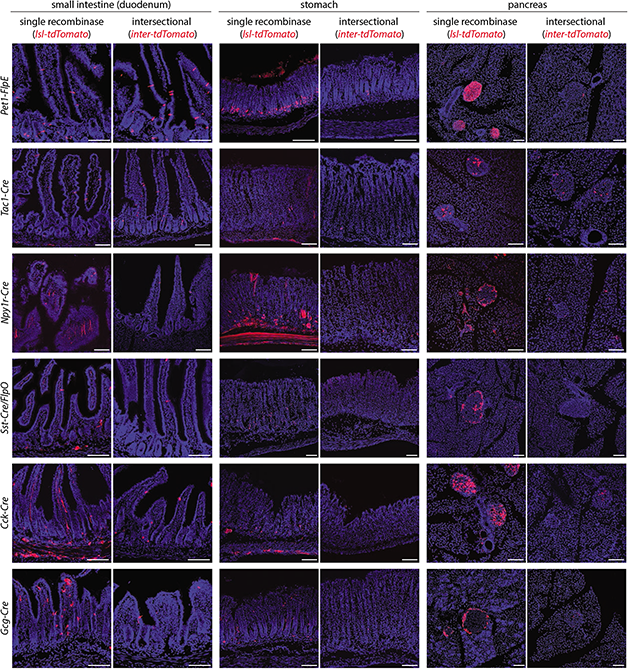
Determining selectivity of tools for intersectional genetics. Native tdTomato fluorescence was analyzed in cryosections (20 μm) of fixed tissues indicated. Tissue was obtained from mice expressing a single recombinase (single recombinase; Cre lines indicated including *Sst-ires-Cre* were crossed with *lsl-tdTomato,* while *Pet1-FlpE* was crossed with *fsf-Gfp* and images pseudocolored in red) or two recombinases (intersectional; Cre lines indicated crossed with *Vil1-p2a-FlpO* and *inter-tdTomato* while *Pet1-FlpE* and *Sst-ires-FlpO* were crossed with *Vil1-Cre* and *inter-tdTomato*). For *Gcg-Cre* mice, images are taken from the ileum rather than duodenum, scale bars: 100 μm.

**Figure 3 - figure supplement 2.**
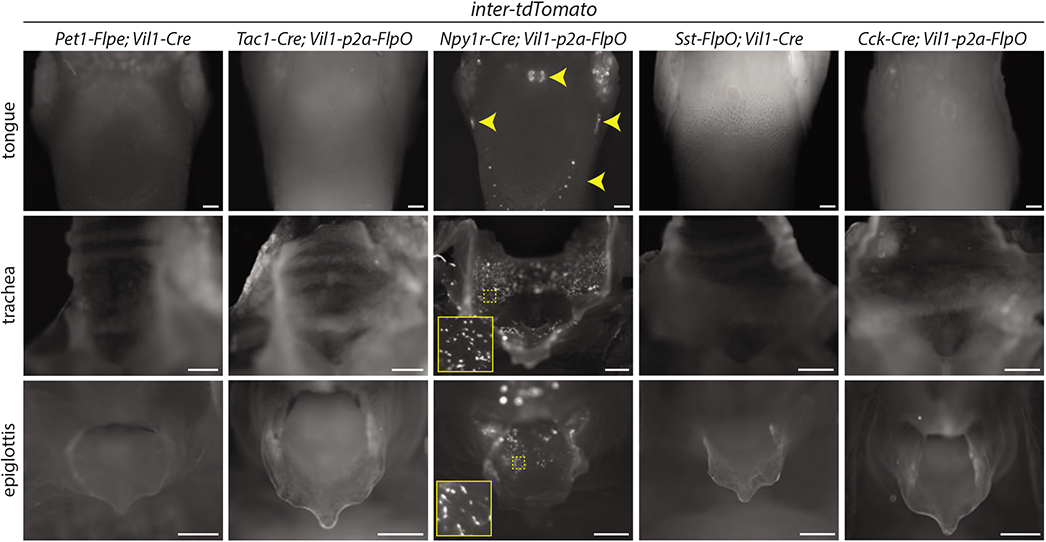
Characterization of reporter expression in the oral cavity and airways. Mouse lines indicated were crossed to *inter-tdTomato* and native reporter fluorescence was analyzed in wholemount tissue preparations from fixative-perfused mice, scale bars: 500 μm.

**Figure 3 - figure supplement 3.**
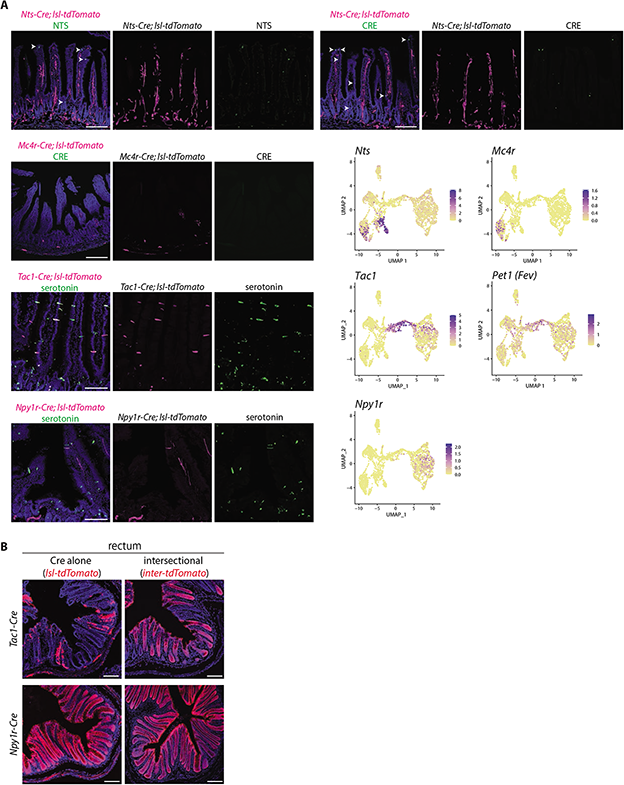
Characterization of genetic tools. (A) Two-color analysis examining expression of tdTomato (native fluorescence, magenta) and either neurotensin (NTS), CRE, or serotonin (immunochemistry, green) in fixed intestinal cryosections (20 μm) of mouse lines indicated. UMAP plots (bottom right) indicate gene expression in single cell RNA sequencing data from enteroendocrine cells. (B) Native tdTomato fluorescence in fixed cryosections from the rectum of mouse lines indicated, scale bars: 100 μm.

**Figure 4 - figure supplement 1.**
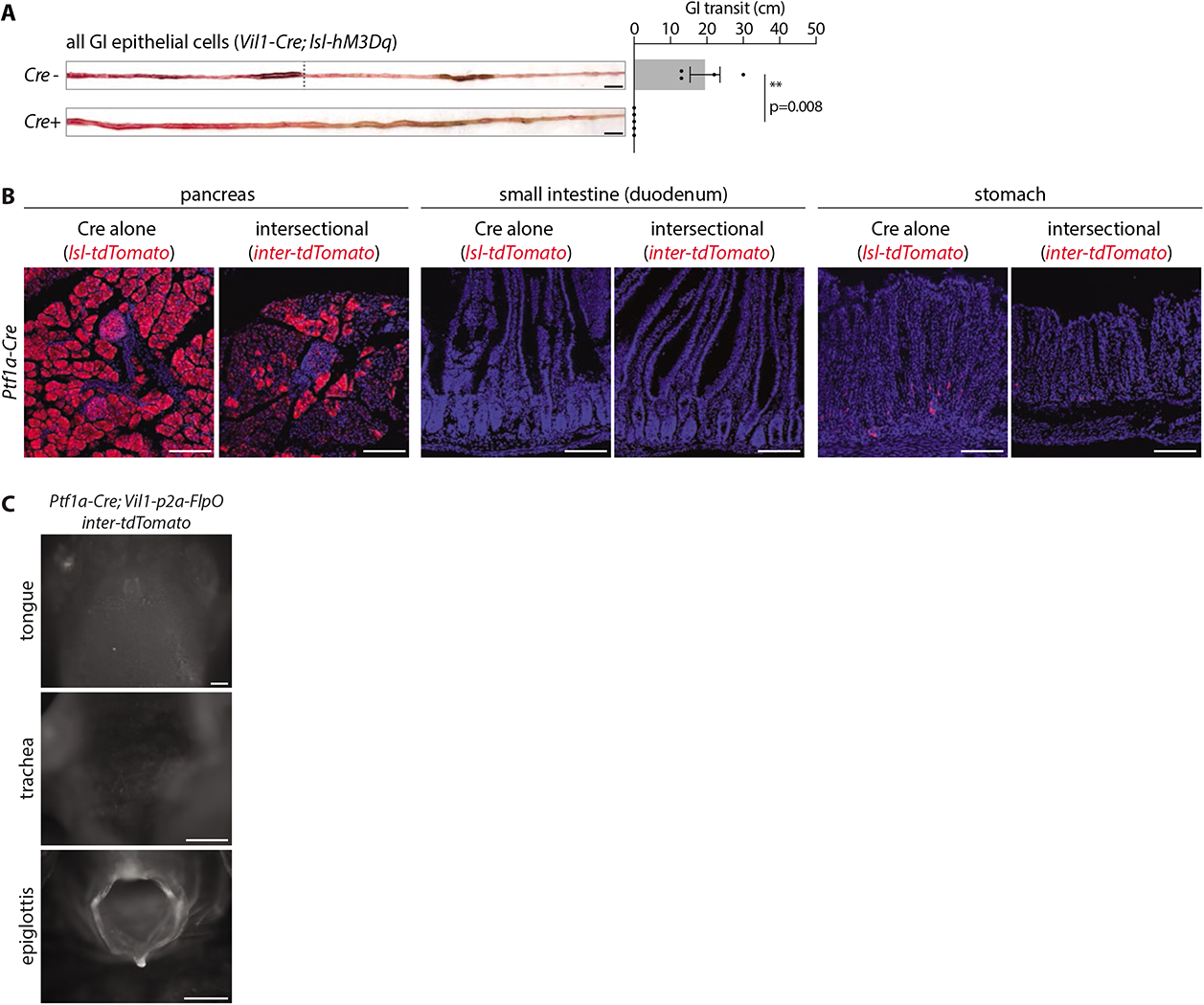
Supporting data for gut transit measurements. (A) Mice indicated were injected with CNO (IP, 3 mg/kg), and gavaged orally with carmen red dye. Intestinal tissue was harvested and the distance between the pyloric sphincter and the charcoal dye leading edge measured. Representative images (left) and quantification (right) of gut transit, scale bar: 1 cm, circles: individual mice, n: 4-5 mice, mean ± sem, **p<0.01 by a Mann-Whitney test. (B) Native tdTomato fluorescence in fixed cryosections (20 μm) of tissues from mice indicated, scale bars: 100 μm. (C) Native reporter fluorescence was analyzed in wholemount tissue preparations indicated, scale bar: 500 μm.

**Figure 5 - figure supplement 1.**
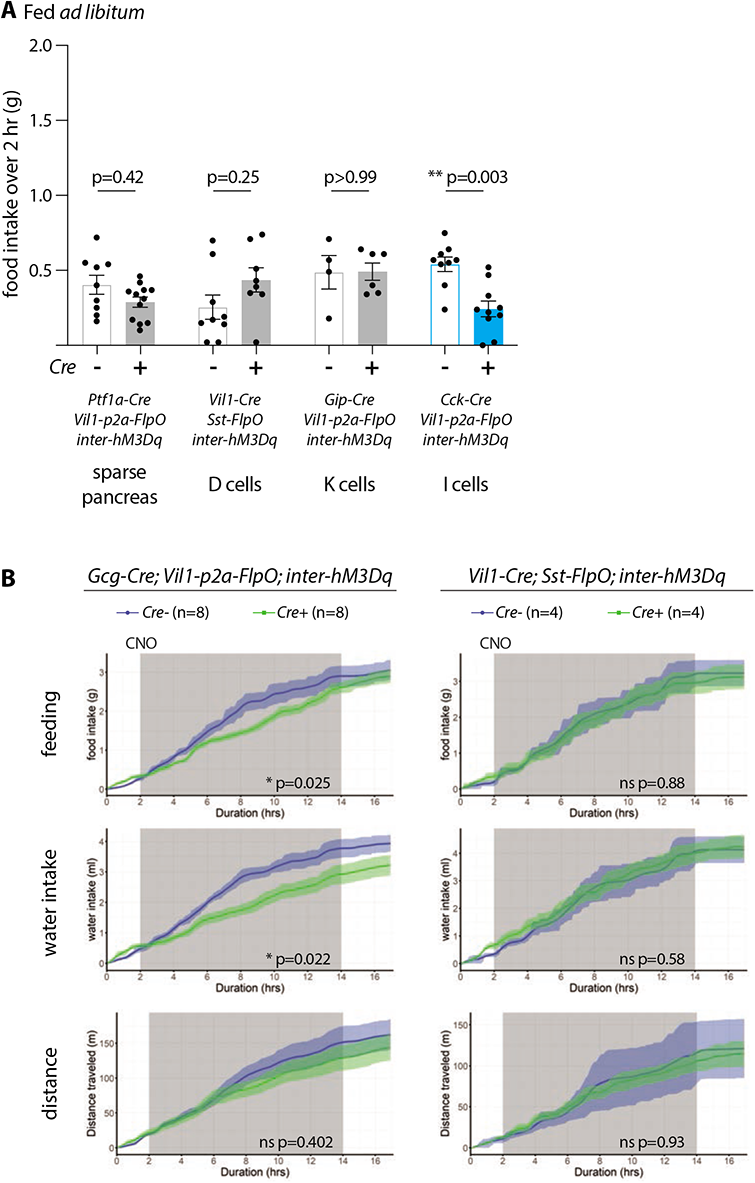
Behavioral responses to enteroendocrine cell activation. (A) Effects of enteroendocrine cell activation on feeding behavior in *ad libitum* fed mice. Mice of genotypes indicated were injected with CNO (IP, 3 mg/kg), and total food intake was measured over two hours, circles: individual mice, n: 4-12 mice, mean ± sem, **p<0.01 by a Mann-Whitney tests with Holm-Šídák correction. (B) Analysis of CNO-evoked behavioral changes in genotypes indicated by a Comprehensive Lab Animal Monitoring System (CLAMS). CNO was injected 15 minutes prior to dark onset (dark: gray shading), n: 4-8 mice, mean ± sem, *p<0.05 by an ANCOVA (feeding, water intake) or ANOVA (distance traveled).

**Figure 6 - figure supplement 1.**
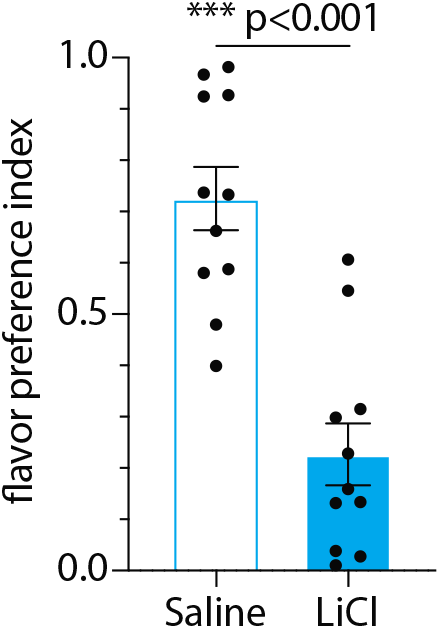
Lithium chloride-induced flavor avoidance. C57BL/6 mice were analyzed for LiCl-evoked flavor avoidance, n=11, mean ± sem, circles: individual mice, ***p<0.001 by a Mann-Whitney test.

## REFERENCES

Alcaino, C., Knutson, K.R., Treichel, A.J., Yildiz, G., Strege, P.R., Linden, D.R., Li, J.H., Leiter, A.B., Szurszewski, J.H., Farrugia, G., and Beyder, A. (2018). A population of gut epithelial enterochromaffin cells is mechanosensitive and requires Piezo2 to convert force into serotonin release. Proc Natl Acad Sci U S A 115, E7632–E7641. 10.1073/pnas.1804938115.

Andermann, M.L., and Lowell, B.B. (2017). Toward a Wiring Diagram Understanding of Appetite Control. Neuron 95, 757–778. 10.1016/j.neuron.2017.06.014.

Baggio, L.L., and Drucker, D.J. (2007). Biology of incretins: GLP-1 and GIP. Gastroenterology 132, 2131–2157. 10.1053/j.gastro.2007.03.054.

Bai, L., Mesgarzadeh, S., Ramesh, K.S., Huey, E.L., Liu, Y., Gray, L.A., Aitken, T.J., Chen, Y., Beutler, L.R., Ahn, J.S., et al. (2019). Genetic Identification of Vagal Sensory Neurons That Control Feeding. Cell 179, 1129–1143 e1123. 10.1016/j.cell.2019.10.031.

Bellono, N.W., Bayrer, J.R., Leitch, D.B., Castro, J., Zhang, C., O’Donnell, T.A., Brierley, S.M., Ingraham, H.A., and Julius, D. (2017). Enterochromaffin Cells Are Gut Chemosensors that Couple to Sensory Neural Pathways. Cell 170, 185–198 e116. 10.1016/j.cell.2017.05.034.

Beumer, J., Artegiani, B., Post, Y., Reimann, F., Gribble, F., Nguyen, T.N., Zeng, H., Van den Born, M., Van Es, J.H., and Clevers, H. (2018). Enteroendocrine cells switch hormone expression along the crypt-to-villus BMP signalling gradient. Nat Cell Biol 20, 909–916. 10.1038/s41556-018-0143-y.

Beumer, J., Gehart, H., and Clevers, H. (2020). Enteroendocrine Dynamics-New Tools Reveal Hormonal Plasticity in the Gut. Endocr Rev 41. 10.1210/endrev/bnaa018.

Beutler, L.R., Chen, Y., Ahn, J.S., Lin, Y.C., Essner, R.A., and Knight, Z.A. (2017). Dynamics of Gut-Brain Communication Underlying Hunger. Neuron 96, 461–475 e465. 10.1016/j.neuron.2017.09.043.

Bohorquez, D.V., Shahid, R.A., Erdmann, A., Kreger, A.M., Wang, Y., Calakos, N., Wang, F., and Liddle, R.A. (2015). Neuroepithelial circuit formed by innervation of sensory enteroendocrine cells. J Clin Invest 125, 782–786. 10.1172/JCI78361.

Brookes, S.J., Spencer, N.J., Costa, M., and Zagorodnyuk, V.P. (2013). Extrinsic primary afferent signalling in the gut. Nat Rev Gastroenterol Hepatol 10, 286–296. 10.1038/nrgastro.2013.29.

Cho, J.H., and Tsai, M.J. (2004). The role of BETA2/NeuroD1 in the development of the nervous system. Mol Neurobiol 30, 35–47. 10.1385/MN:30:1:035.

de Araujo, I.E., Oliveira-Maia, A.J., Sotnikova, T.D., Gainetdinov, R.R., Caron, M.G., Nicolelis, M.A., and Simon, S.A. (2008). Food reward in the absence of taste receptor signaling. Neuron 57, 930–941. 10.1016/j.neuron.2008.01.032.

Drucker, D.J. (2016). Evolving Concepts and Translational Relevance of Enteroendocrine Cell Biology. J Clin Endocrinol Metab 101, 778–786. 10.1210/jc.2015-3449.

el Marjou, F., Janssen, K.P., Chang, B.H., Li, M., Hindie, V., Chan, L., Louvard, D., Chambon, P., Metzger, D., and Robine, S. (2004). Tissue-specific and inducible Cre-mediated recombination in the gut epithelium. Genesis 39, 186–193. 10.1002/gene.20042.

Fernandes, A.B., Alves da Silva, J., Almeida, J., Cui, G., Gerfen, C.R., Costa, R.M., and Oliveira-Maia, A.J. (2020). Postingestive Modulation of Food Seeking Depends on Vagus-Mediated Dopamine Neuron Activity. Neuron 106, 778–788 e776. 10.1016/j.neuron.2020.03.009.

Freeman, A.J., Cunningham, K.T., and Tyers, M.B. (1992). Selectivity of 5-HT3 receptor antagonists and anti-emetic mechanisms of action. Anticancer Drugs 3, 79–85.

Gale, J.D. (1995). Serotonergic mediation of vomiting. J Pediatr Gastroenterol Nutr 21 *Suppl 1*, S22–28. 10.1097/00005176-199501001-00008.

Gao, Z., Ure, K., Ables, J.L., Lagace, D.C., Nave, K.A., Goebbels, S., Eisch, A.J., and Hsieh, J. (2009). Neurod1 is essential for the survival and maturation of adult-born neurons. Nat Neurosci 12, 1090–1092. 10.1038/nn.2385.

Gehart, H., van Es, J.H., Hamer, K., Beumer, J., Kretzschmar, K., Dekkers, J.F., Rios, A., and Clevers, H. (2019). Identification of Enteroendocrine Regulators by Real-Time Single-Cell Differentiation Mapping. Cell 176, 1158–1173 e1116. 10.1016/j.cell.2018.12.029.

Gorboulev, V., Schurmann, A., Vallon, V., Kipp, H., Jaschke, A., Klessen, D., Friedrich, A., Scherneck, S., Rieg, T., Cunard, R., et al. (2012). Na(+)-D-glucose cotransporter SGLT1 is pivotal for intestinal glucose absorption and glucose-dependent incretin secretion. Diabetes 61, 187–196. 10.2337/db11-1029.

Gribble, F.M., and Reimann, F. (2019). Function and mechanisms of enteroendocrine cells and gut hormones in metabolism. Nat Rev Endocrinol 15, 226–237. 10.1038/s41574-019-0168-8.

Gribble, F.M., and Reimann, F. (2021). Metabolic Messengers: glucagon-like peptide 1. Nat Metab 3, 142–148. 10.1038/s42255-020-00327-x.

Haber, A.L., Biton, M., Rogel, N., Herbst, R.H., Shekhar, K., Smillie, C., Burgin, G., Delorey, T.M., Howitt, M.R., Katz, Y., et al. (2017). A single-cell survey of the small intestinal epithelium. Nature 551, 333–339. 10.1038/nature24489.

Habib, A.M., Richards, P., Cairns, L.S., Rogers, G.J., Bannon, C.A., Parker, H.E., Morley, T.C., Yeo, G.S., Reimann, F., and Gribble, F.M. (2012). Overlap of endocrine hormone expression in the mouse intestine revealed by transcriptional profiling and flow cytometry. Endocrinology 153, 3054–3065. 10.1210/en.2011-2170.

Hafemeister, C., and Satija, R. (2019). Normalization and variance stabilization of single-cell RNA-seq data using regularized negative binomial regression. Genome Biol 20, 296. 10.1186/s13059-019-1874-1.

Han, W., Tellez, L.A., Perkins, M.H., Perez, I.O., Qu, T., Ferreira, J., Ferreira, T.L., Quinn, D., Liu, Z.W., Gao, X.B., et al. (2018). A Neural Circuit for Gut-Induced Reward. Cell 175, 887–888. 10.1016/j.cell.2018.10.018.

Hofer, D., and Drenckhahn, D. (1999). Localisation of actin, villin, fimbrin, ezrin and ankyrin in rat taste receptor cells. Histochem Cell Biol 112, 79–86. 10.1007/s004180050394.

Holst, J.J., Vilsboll, T., and Deacon, C.F. (2009). The incretin system and its role in type 2 diabetes mellitus. Mol Cell Endocrinol 297, 127–136. 10.1016/j.mce.2008.08.012.

Jenny, M., Uhl, C., Roche, C., Duluc, I., Guillermin, V., Guillemot, F., Jensen, J., Kedinger, M., and Gradwohl, G. (2002). Neurogenin3 is differentially required for endocrine cell fate specification in the intestinal and gastric epithelium. EMBO J 21, 6338–6347. 10.1093/emboj/cdf649.

Jensen, P., Farago, A.F., Awatramani, R.B., Scott, M.M., Deneris, E.S., and Dymecki, S.M. (2008). Redefining the serotonergic system by genetic lineage. Nat Neurosci 11, 417–419. 10.1038/nn2050.

Kawaguchi, Y., Cooper, B., Gannon, M., Ray, M., MacDonald, R.J., and Wright, C.V. (2002). The role of the transcriptional regulator Ptf1a in converting intestinal to pancreatic progenitors. Nat Genet 32, 128–134. 10.1038/ng959.

Lee, S.Y., and Soltesz, I. (2011). Cholecystokinin: a multi-functional molecular switch of neuronal circuits. Dev Neurobiol 71, 83–91. 10.1002/dneu.20815.

Lewis, J.E., Miedzybrodzka, E.L., Foreman, R.E., Woodward, O.R.M., Kay, R.G., Goldspink, D.A., Gribble, F.M., and Reimann, F. (2020). Selective stimulation of colonic L cells improves metabolic outcomes in mice. Diabetologia 63, 1396–1407. 10.1007/s00125-020-05149-w.

Li, H.J., Kapoor, A., Giel-Moloney, M., Rindi, G., and Leiter, A.B. (2012). Notch signaling differentially regulates the cell fate of early endocrine precursor cells and their maturing descendants in the mouse pancreas and intestine. Dev Biol 371, 156–169. 10.1016/j.ydbio.2012.08.023.

Li, H.J., Ray, S.K., Singh, N.K., Johnston, B., and Leiter, A.B. (2011). Basic helix-loop-helix transcription factors and enteroendocrine cell differentiation. Diabetes Obes Metab 13 *Suppl 1*, 5–12. 10.1111/j.1463-1326.2011.01438.x.

Madison, B.B., Dunbar, L., Qiao, X.T., Braunstein, K., Braunstein, E., and Gumucio, D.L. (2002). Cis elements of the villin gene control expression in restricted domains of the vertical (crypt) and horizontal (duodenum, cecum) axes of the intestine. J Biol Chem 277, 33275–33283. 10.1074/jbc.M204935200.

Maunoury, R., Robine, S., Pringault, E., Leonard, N., Gaillard, J.A., and Louvard, D. (1992). Developmental regulation of villin gene expression in the epithelial cell lineages of mouse digestive and urogenital tracts. Development 115, 717–728.

Mellitzer, G., Beucher, A., Lobstein, V., Michel, P., Robine, S., Kedinger, M., and Gradwohl, G. (2010). Loss of enteroendocrine cells in mice alters lipid absorption and glucose homeostasis and impairs postnatal survival. J Clin Invest 120, 1708–1721. 10.1172/JCI40794.

Mina, A.I., LeClair, R.A., LeClair, K.B., Cohen, D.E., Lantier, L., and Banks, A.S. (2018). CalR: A Web-Based Analysis Tool for Indirect Calorimetry Experiments. Cell Metab 28, 656–666 e651. 10.1016/j.cmet.2018.06.019.

Naya, F.J., Huang, H.P., Qiu, Y., Mutoh, H., DeMayo, F.J., Leiter, A.B., and Tsai, M.J. (1997). Diabetes, defective pancreatic morphogenesis, and abnormal enteroendocrine differentiation in BETA2/neuroD-deficient mice. Genes Dev 11, 2323–2334. 10.1101/gad.11.18.2323.

Nozawa, K., Kawabata-Shoda, E., Doihara, H., Kojima, R., Okada, H., Mochizuki, S., Sano, Y., Inamura, K., Matsushime, H., Koizumi, T., et al. (2009). TRPA1 regulates gastrointestinal motility through serotonin release from enterochromaffin cells. Proc Natl Acad Sci U S A 106, 3408–3413. 10.1073/pnas.0805323106.

Okaty, B.W., Commons, K.G., and Dymecki, S.M. (2019). Embracing diversity in the 5-HT neuronal system. Nat Rev Neurosci 20, 397–424. 10.1038/s41583-019-0151-3.

Prescott, S.L., and Liberles, S.D. (2022). Internal senses of the vagus nerve. Neuron. 10.1016/j.neuron.2021.12.020.

Reimann, F., Habib, A.M., Tolhurst, G., Parker, H.E., Rogers, G.J., and Gribble, F.M. (2008). Glucose sensing in L cells: a primary cell study. Cell Metab 8, 532–539. 10.1016/j.cmet.2008.11.002.

Reimann, F., Tolhurst, G., and Gribble, F.M. (2012). G-protein-coupled receptors in intestinal chemosensation. Cell Metab 15, 421–431. 10.1016/j.cmet.2011.12.019.

Richards, P., Thornberry, N.A., and Pinto, S. (2021). The gut-brain axis: Identifying new therapeutic approaches for type 2 diabetes, obesity, and related disorders. Mol Metab 46, 101175. 10.1016/j.molmet.2021.101175.

Roth, B.L. (2016). DREADDs for Neuroscientists. Neuron 89, 683–694. 10.1016/j.neuron.2016.01.040.

Rutlin, M., Rastelli, D., Kuo, W.T., Estep, J.A., Louis, A., Riccomagno, M.M., Turner, J.R., and Rao, M. (2020). The Villin1 Gene Promoter Drives Cre Recombinase Expression in Extraintestinal Tissues. Cell Mol Gastroenterol Hepatol 10, 864–867 e865. 10.1016/j.jcmgh.2020.05.009.

Satija, R., Farrell, J.A., Gennert, D., Schier, A.F., and Regev, A. (2015). Spatial reconstruction of single-cell gene expression data. Nat Biotechnol 33, 495–502. 10.1038/nbt.3192.

Sato, T., Vries, R.G., Snippert, H.J., van de Wetering, M., Barker, N., Stange, D.E., van Es, J.H., Abo, A., Kujala, P., Peters, P.J., and Clevers, H. (2009). Single Lgr5 stem cells build crypt-villus structures in vitro without a mesenchymal niche. Nature 459, 262–265. 10.1038/nature07935.

Schonhoff, S.E., Giel-Moloney, M., and Leiter, A.B. (2004). Neurogenin 3-expressing progenitor cells in the gastrointestinal tract differentiate into both endocrine and non-endocrine cell types. Dev Biol 270, 443–454. 10.1016/j.ydbio.2004.03.013.

Sciolino, N.R., Plummer, N.W., Chen, Y.W., Alexander, G.M., Robertson, S.D., Dudek, S.M., McElligott, Z.A., and Jensen, P. (2016). Recombinase-Dependent Mouse Lines for Chemogenetic Activation of Genetically Defined Cell Types. Cell Rep 15, 2563–2573. 10.1016/j.celrep.2016.05.034.

Sclafani, A., Koepsell, H., and Ackroff, K. (2016). SGLT1 sugar transporter/sensor is required for post-oral glucose appetition. Am J Physiol Regul Integr Comp Physiol 310, R631–639. 10.1152/ajpregu.00432.2015.

Seeley, R.J., Chambers, A.P., and Sandoval, D.A. (2015). The role of gut adaptation in the potent effects of multiple bariatric surgeries on obesity and diabetes. Cell Metab 21, 369–378. 10.1016/j.cmet.2015.01.001.

Slotnick, B. (2009). A simple 2-transistor touch or lick detector circuit. J Exp Anal Behav 91, 253–255. 10.1901/jeab.2009.91-253.

Sternson, S.M., and Eiselt, A.K. (2017). Three Pillars for the Neural Control of Appetite. Annu Rev Physiol 79, 401–423. 10.1146/annurev-physiol-021115-104948.

Stuart, T., Butler, A., Hoffman, P., Hafemeister, C., Papalexi, E., Mauck, W.M., 3rd, Hao, Y., Stoeckius, M., Smibert, P., and Satija, R. (2019). Comprehensive Integration of Single-Cell Data. Cell 177, 1888–1902 e1821. 10.1016/j.cell.2019.05.031.

Tan, H.E., Sisti, A.C., Jin, H., Vignovich, M., Villavicencio, M., Tsang, K.S., Goffer, Y., and Zuker, C.S. (2020). The gut-brain axis mediates sugar preference. Nature 580, 511–516. 10.1038/s41586-020-2199-7.

Treichel, A.J., Finholm, I., Knutson, K.R., Alcaino, C., Whiteman, S.T., Brown, M.R., Matveyenko, A., Wegner, A., Kacmaz, H., Mercado-Perez, A., et al. (2022). Specialized Mechanosensory Epithelial Cells in Mouse Gut Intrinsic Tactile Sensitivity. Gastroenterology 162, 535–547 e513. 10.1053/j.gastro.2021.10.026.

Tyers, M.B., and Freeman, A.J. (1992). Mechanism of the anti-emetic activity of 5-HT3 receptor antagonists. Oncology 49, 263–268. 10.1159/000227054.

Van Citters, G.W., and Lin, H.C. (2006). Ileal brake: neuropeptidergic control of intestinal transit. Curr Gastroenterol Rep 8, 367–373. 10.1007/s11894-006-0021-9.

Wang, J., Cortina, G., Wu, S.V., Tran, R., Cho, J.H., Tsai, M.J., Bailey, T.J., Jamrich, M., Ament, M.E., Treem, W.R., et al. (2006). Mutant neurogenin-3 in congenital malabsorptive diarrhea. N Engl J Med 355, 270–280. 10.1056/NEJMoa054288.

Williams, E.K., Chang, R.B., Strochlic, D.E., Umans, B.D., Lowell, B.B., and Liberles, S.D. (2016). Sensory Neurons that Detect Stretch and Nutrients in the Digestive System. Cell 166, 209–221. 10.1016/j.cell.2016.05.011.

Yang, H., Xie, X., Deng, M., Chen, X., and Gan, L. (2010). Generation and characterization of Atoh1-Cre knock-in mouse line. Genesis 48, 407–413. 10.1002/dvg.20633.

Yarmolinsky, D.A., Zuker, C.S., and Ryba, N.J. (2009). Common sense about taste: from mammals to insects. Cell 139, 234–244. 10.1016/j.cell.2009.10.001.

Zhang, C., Kaye, J.A., Cai, Z., Wang, Y., Prescott, S.L., and Liberles, S.D. (2021). Area Postrema Cell Types that Mediate Nausea-Associated Behaviors. Neuron 109, 461–472 e465. 10.1016/j.neuron.2020.11.010.

Zimmerman, C.A., and Knight, Z.A. (2020). Layers of signals that regulate appetite. Curr Opin Neurobiol 64, 79–88. 10.1016/j.conb.2020.03.007.

